# Cell region fingerprints enable highly precise single-cell tracking and lineage reconstruction

**DOI:** 10.1101/2021.10.26.465883

**Authors:** Andreas P. Cuny, Aaron Ponti, Tomas Kündig, Fabian Rudolf, Jörg Stelling

**Affiliations:** Department of Biosystems Science and Engineering, ETH Zurich, Mattenstrasse 26, 4058 Basel; Swiss Institute of Bioinformatics, Mattenstrasse 26, 4058 Basel

## Abstract

Experimental studies of cell growth, inheritance, and their associated processes by microscopy require accurate single-cell observations of sufficient duration to reconstruct the genealogy. However, cell tracking—assigning identical cells on consecutive images to a track—is often challenging due to imperfect segmentation, moving cells, or focus drift, resulting in laborious manual verification. Here, we propose fingerprints to identify problematic assignments rapidly. A fingerprint distance measures the similarity between cells in two consecutive images by comparing the structural information contained in the low frequencies of a Fourier transform. We show that it is broadly applicable across cell types and image modalities, provided the image has sufficient structural information. Our tracker (*Trac*^*X*^) uses the concept to reject unlikely assignments, thereby substantially increasing tracking performance on published and newly generated long-term data sets from various species. For *S. cerevisiae*, we propose a comprehensive model for cell size control at the single-cell and population level centered on the Whi5 regulator. It demonstrates how highly precise tracking can help uncover previously undescribed single-cell biology.

## Introduction

Live-cell imaging is the primary tool to study dynamics processes and their inheritance across generations at the single-cell level. It also enables novel applications to identify and isolate cells with a dynamic phenotype of interest^1^ and to connect genotype with phenotype by imaging-based screening^2^. The analysis starts from often large sets of raw images (**Fig. 1a**), in which objects such as cells have to be identified by segmentation methods (**Fig. 1b**). One then needs to establish the temporal relationship between all the segmented objects in each image in a task termed tracking (**Fig. 1c**). Accurate tracking is essential, especially when studying inheritance or differentiation, which requires complete cell lineages, that is, the genealogy (**Fig. 1d**). For example, lineage-based analysis recently revealed long-term memory linking cell growth and cell cycle progression in mammalian cells^3^ and suggested asymmetric inheritance as a key mechanism determining hematopoietic stem cell fate^4^.

**Figure 1.**
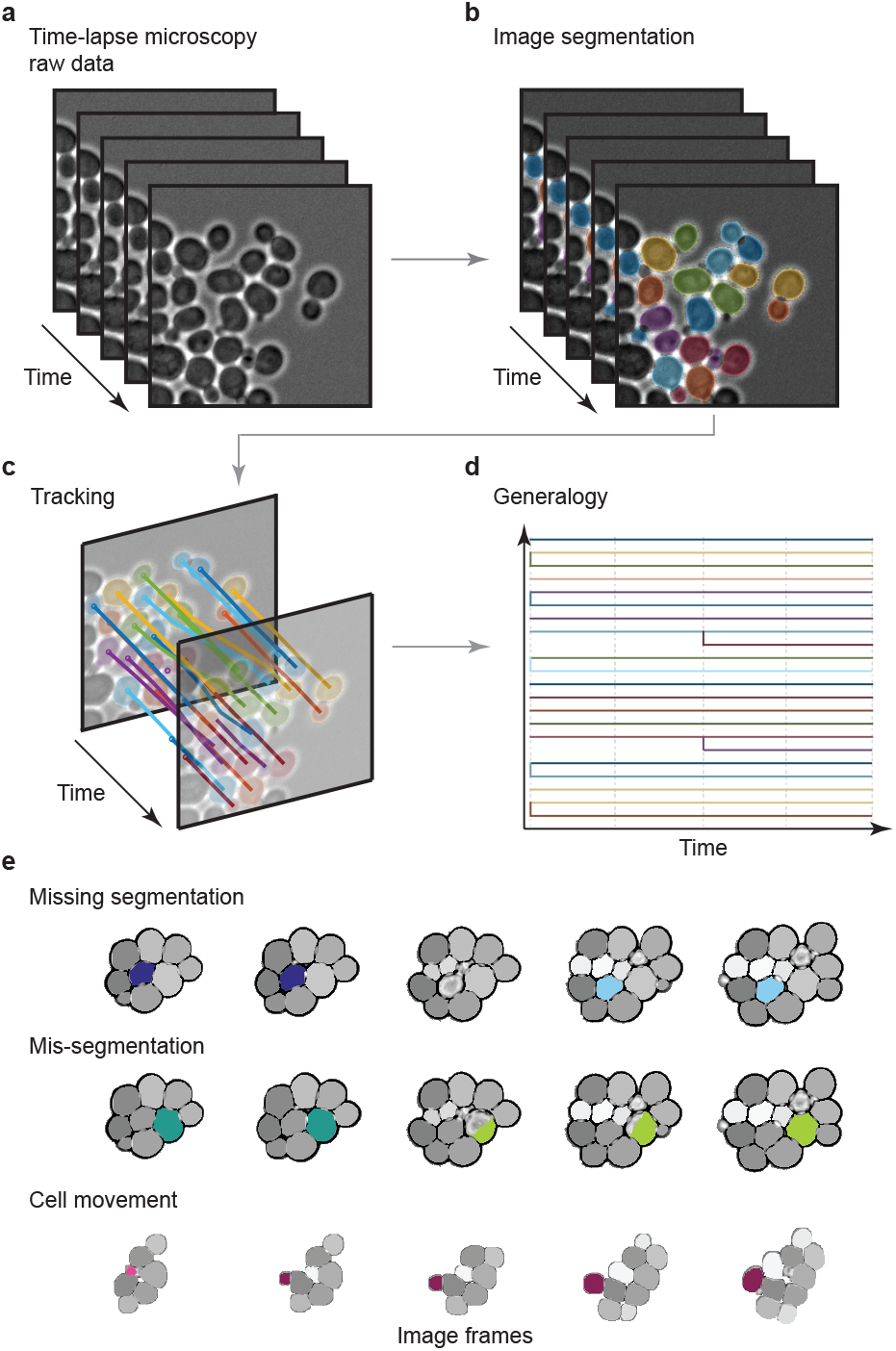
Cell tracking pipeline and common problems. **a** Typical time-lapse microscopy dataset: bright field images from various time-points as input. **b** Image segmentation for the detection of cells (colored). **c** Tracking of segmented objects over (consecutive) time-points. **d** Reconstruction of the genealogy. Colors in **b-d** indicate cell identity. **e** Common difficulties during cell tracking over consecutive frames (columns) illustrated by individual cells (colored) and their track associations (colors). First row: Missing segmentation in frame 3 leads to premature track termination in frame 2, resulting in two separate short tracks (frames 1-2 and 4-5) without assignment in frame 3. Second row: Mis-segmentation in frame 3 leads to the restart of a new track in frame 3, resulting in two short tracks (1-2 and 3-5). Third row: Cell displacement due to cell growth or culturing creates artificial track ends (frame 1) and starts (frame 2).

With important progress in cell segmentation methods that often involve deep learning approaches^5^ and generalize across cell types and imaging modalities^6^, cell tracking is the main current bottleneck for reliable image analysis at high throughput^7^. In addition to inaccurate segmentation results used for tracking, different events during the life of a cell, such as birth, division, changing morphology during growth, death, and migration (see **Fig. 1e** for examples) make tracking a challenging task. Importantly, incorrect assignments of cells to tracks propagate during lineage reconstruction, such that even minor tracking inaccuracies can impact the biological interpretation of the imaging data substantially.

In addressing the well-recognized need for better tracking tools^8–11^, methods for the automated validation of tracking results are critical. They could alleviate the limitations on accuracy and throughput imposed by the need for laborious manual curation. Current tools only allow for a software-assisted or user-driven detection of tracking errors based on a resulting lineage^9^ and for a manual inspection of tracking results^8,11,12^ with possible interactive correction^13–15^. First deep learning-based tracking methods show great promise, for example, with 1% error rate when extensive training data was available, but non-competitive results with limited training data^16^. The problem of manual curation to establish a ground truth for cell tracking, hence, is a general one when large data sets need to be processed for analysis or training purposes. It is even more extreme for real-time applications such as automated control of single-cell behavior using optogenetics^17^.

To assess cell tracking results automatically, we took inspiration from approaches in image watermarking and matching that compare the structures of images^18–20^. We reasoned that we can compare a cell and its surrounding between consecutive images to identify correct cell-to-track assignments. Specifically, we devised a cell region fingerprint (*CRF*) that captures the structural information, that is, basic image attributes representing the structure of depicted objects, and not luminance and contrast. It does not require extensive training because we need few parameters that have a direct physical interpretation and can be estimated from a few image frames. The approach allows to identify problematic tracking results even when the ground truth is unknown and it is independent of, for example, specific segmentation or tracking algorithms. We use the *CRF* in a new tracker that, akin to kinetic proofreading, rejects linkages likely to be wrong. Performance tests of our methods with published data sets and newly acquired data for various cell types and image modalities demonstrate near-perfect tracking accuracy. This allowed us to reconstruct the genealogy of symmetrically and asymmetrically dividing cell types in general, and to analyze *S. cerevisiae*’s cell size regulation and homeostasis comprehensively.

## Results

### Cell region fingerprints capture relevant structural information

A cell region fingerprint (*CRF*) aims to characterize a cell and its surrounding such that we can use these characteristics to identify correct cell-to-track assignments. We calculate the fingerprint from the same raw images used for segmentation, by defining a square region around a cell’s centroid to also include the cell’s neighborhood (**Fig. 2a**; see **Methods** for details). Note that one can use any segmentation tool that creates a mask of the segmented objects (labeled segmentation mask) to derive the cell centroids. Individual badly segmented cells will be detected in a time series by a changing signature, but the *CRF* alone does not tell if a segmentation is good or bad. After cropping the rest of the image (**Fig. 2b**), we scale the image matrix to a defined size that is identical for all cells (**Fig. 2c**). To extract structural information, we Fourier transform the image using the discrete cosine transformation (DCT; **Fig. 2d**). The lowest frequencies of the DCT contain the structural information of the image, as demonstrated by the inverse DCT in **Fig. 2e**. We therefore define the *CRF* as a matrix that contains only DCT coefficients for the lowest frequencies (**Fig. 2f**). This procedure implies that the *CRF* computation depends only on three parameters: the size of the image window, the re-scaling factor, and the number of DCT frequencies to include.

**Figure 2.**
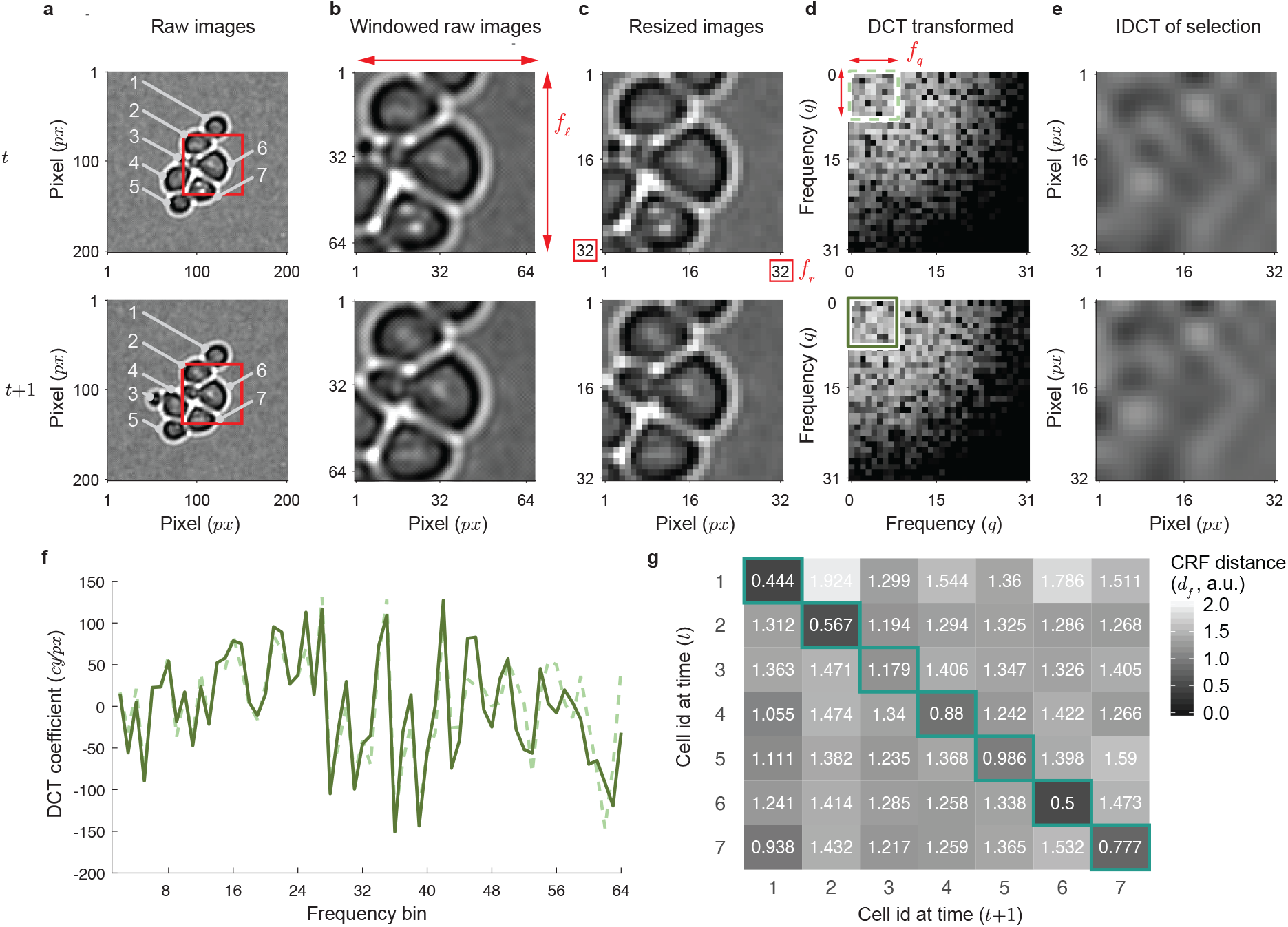
Cell region fingerprint (*CRF*). **a** Raw images of two frames *t* and *t* + 1 of a *S. cerevisiae* cell culture. Numbers: cell identifiers; red squares: cropping regions of approximately twice the cell diameter around the centroid of cell 6. **b** Cropped image regions. **c** Re-sized image crops (32 × 32 pixels). **d** Discrete cosine transformation (DCT) of images in **c**. Green squares: lowest frequencies (8 × 8) used as cell region fingerprint. **e** Inverse DCT (iDCT) of the 8 × 8 frequencies. **f** Superimposed, arrayed 8 × 8 frequencies of the *CRF*. Solid line: frame *t*; dashed line: frame *t* + 1. **g** Cell region fingerprint distance (*d*_*f*_) matrix for all pairs of cells in frames *t* and *t* + 1 from **a**. Green boxes highlight lowest distances between a cell in *t* and any cell in *t* + 1.

To compare any pair of cells in principle, but possible assignments between pairs of cells in two consecutive images in live-cell microscopy for the tracking problem specifically, we next define a fingerprint distance as a scalar measure of similarity. It is the normalized sum of squared residuals between two *CRF* arrays, denoted by *d*_*f*_ (see **Methods**). By construction, *d*_*f*_ = 0 for identical image regions, and it increases the more the two regions differ. **Fig. 2g** indicates that the information condensed in the fingerprint distance may indeed be sufficient to distinguish between correct and incorrect tracking assignments: the correctly assigned cell pairs on the diagonal have the lowest *d*_*f*_ values of all possible assignments.

To construct a classifier that distinguishes between correct and incorrect tracking results, we reasoned that the *CRF* is less likely to be unique for an entire microscopy image because geometrically simple cells such as round or rod-shaped cells show repeating patterns within a densely populated image. We therefore limit the comparison of fingerprints via the fingerprint distance to smaller regions, namely the neighborhood of a cell of interest. Specifically, we compute *d*_*f*_ between a cell in the preceding frame and the assigned cell in the current frame as well as this cell’s neighbors. Our measure for classification is then the fraction of neighboring cells in the current frame that has a lower *d*_*f*_ than the assigned cell, the neighborhood fraction denoted by *F*_*f*_ (see **Methods** for details). We reason that higher values of *F*_*f*_ suggest more possible alternative assignments, and thereby a higher likelihood that the given assignment is incorrect. Overall, thus, we condense the structural information of cells and their surrounding into a simple and intuitive measure for the evaluation of tracking results for each cell.

### Fingerprints classify tracking assignments reliably and generically

A classifier for tracking assignments needs to reliably distinguish correct from incorrect assignments. It should also be generic (applying to multiple cell types and imaging modalities) and robust (insensitive to specific parameter settings and experimental artifacts). To assess these aspects of *CRF*-based classification, we obtained or constructed the ground truth for published data sets of different cell types, image modalities, and magnifications (see examples in **Fig. 3a** and **Methods**). We then randomly permuted 1% of the assignments to create faulty tracks. Neighborhood fractions (*F*_*f*_) were computed with (jointly) varying parameter values for window side length (*f*_*l*_), re-size factor (*f*_*r*_), and number of DCT coefficients (*f*_*q*_). Finally, we classified the assignments by whether *F*_*f*_ exceeds a fixed, global threshold *τ*_*f*_. Specifically, we used *F*_*f*_ > *τ*_*f*_ (*F*_*f*_ ≤ *τ*_*f*_) to predict incorrect (correct) assignments (see **Methods**).

**Figure 3.**
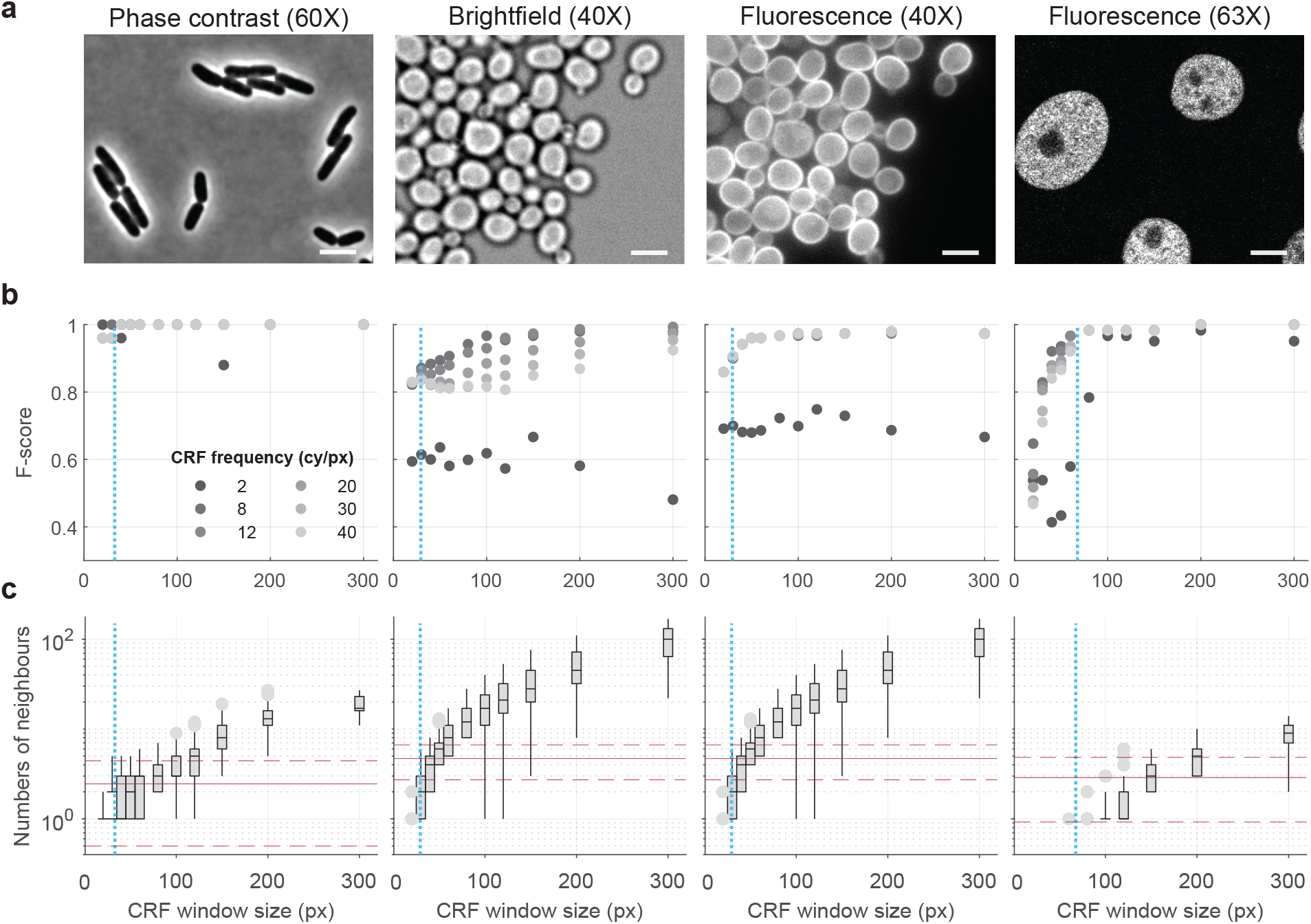
Cell region fingerprint parameter evaluation for different cell types, image modalities and magnifications. **a** Example images for bacterial (*E. coli*^48^; left), yeast (*S. cerevisiae*; middle) and mammalian (N2DH-GOWT1 mouse embryonic stem cells;^33^ right) cells acquired as indicated. Scale bar: 32 pixels. **b** F-scores depending on CRF window sizes (*f*_*l*_) and CRF frequencies (*f*_*q*_) for a fixed resizing factor (*f*_*r*_ =121 px or 4.66 ± 0.05 neighbouring cells; the value did not influence the F-score, see **Fig. S1**). Dotted cyan lines: mean cell diameters. **c**. Numbers of neighbour cells depending on window size (Box plots). Numbers of neighbours for computing the fraction (*F*_*f*_) are given as mean (red solid line) and 95% confidence interval (red dashed lines).

We conducted systematic performance tests by evaluating predictions against the ground truth, initially with a conservative *τ*_*f*_ = 0. Using the F-score (the harmonic mean of precision and recall; see **Methods**) as performance metric revealed five features of our *CRF*-based classifier. First, it can achieve near-perfect performance for all our test cases, which include different cell types (bacterial, yeast, and mammalian) as well as imaging modalities (phase contrast, bright field, and fluorescence; **Fig. 3b**). Second, to capture enough structural information in the image, a low number of DCT coefficients (*f*_*q*_ ≈ 10) is optimal. Third and interestingly, the classifier performance increases monotonically with window size in the tested range (**Fig. 3b**), although the corresponding neighborhoods can include a hundred cells (**Fig. 3c**). Fourth, across all test cases, performance is consistently high for common parameter settings of *f*_*q*_ = 8 and window size of twice the mean cell diameter (corresponding to ≈2-10 neighboring cells; **Fig. 3b,c**). Finally, classifier performance does not depend on the re-size factor (*f*_*r*_; **Fig. S1**).

Overall, our *CRF*-based classifier shows robust and generic high performance. Reliable classification requires only generic parameters, or parameters easily derived from the images, such as the cell diameter. Importantly, setting the threshold of a classifier without knowing the ground truth is a frequent problem—but as additional tests for all image modalities showed (**Fig. S2**), not for the *CRF*-based evaluation of tracking assignments.

### Assignment proof reading improves tracker performance

To demonstrate practical relevance of our classifier for live cell tracking, we aimed to improve an existing tracker to be able to handle imperfect input, such as missing or miss-segmentation and moving cells. The basic idea was to proofread (preliminary) assignments with the *CRF*-based classifier and to re-evaluate questionable ones. We termed the tracker *Trac*^*X*^, analogous to our segmentation software *CellX*^21,22^.

Briefly, we adapted a simple frame-by-frame tracker^23^ to obtain raw assignments. We use cell features such as position, size, and orientation to construct a cost matrix and then solve the linear assignment problem (LAP) by minimizing the cost matrix between two consecutive frames with the Jonker-Volgenant algorithm^24^, an improved version of Kuhn’s algorithm^25^. This yields assigned and non-assigned cells. Next, we classify the raw assignments. Those that do not pass the classifier are deleted and the corresponding cells are designated non-assigned. All non-assigned cell are then joined (for a certain ‘lifetime’) with all cells in the next frame(s) for re-evaluation. However, this approach only works for cells with a slowly changing neighborhood. To capture when cells move rapidly or two growing colonies fuse, we built on the idea of neighborhood preserved motion: the motion of a single cell needs to follow the motion of its direct neighborhood^26^. Our tracking refinement module (**Fig. S7)** uses the assignments that pass the classifier to form a temporary ground truth. From it, we estimate the vector field of cell movement for the re-evaluation of assignments (see **Methods** and **Supplementary Text** for details).

To evaluate the performance of *Trac*^*X*^ systematically in comparison to state-of-the-art trackers, we used the yeast image toolkit (YIT)^11^. It is a collection of manually curated *S. cerevisiae* imaging data sets with different numbers of cells and image frames, designed for systematic comparisons of segmentation and tracking tools. Specifically, we compared the long-term tracking performance of *Trac*^*X*^ measured by an F-score (see **Methods** for details) to the published results for the five best-performing trackers on these data sets^10,11^: CellTracer^27^, CellProfiler^28^, CellID^29^, CellStar^11^, and the algorithm by Wood *et al*.^30^. The comparatively simple *Trac*^*X*^ tracked all YIT data sets, and also quantitatively it outperformed all the other algorithms (**Fig. 4a**). A tracker based on the fingerprint distance alone (*Trac*^*X*^_*CRF*_) did not yield consistent long-term performance. Data set YIT-TS3 stands out: only our algorithm assigns all cells correctly in this time series of densely growing cells with one colony translating and merging with two other colonies (exemplified in **Fig. 4b**).

**Figure 4.**
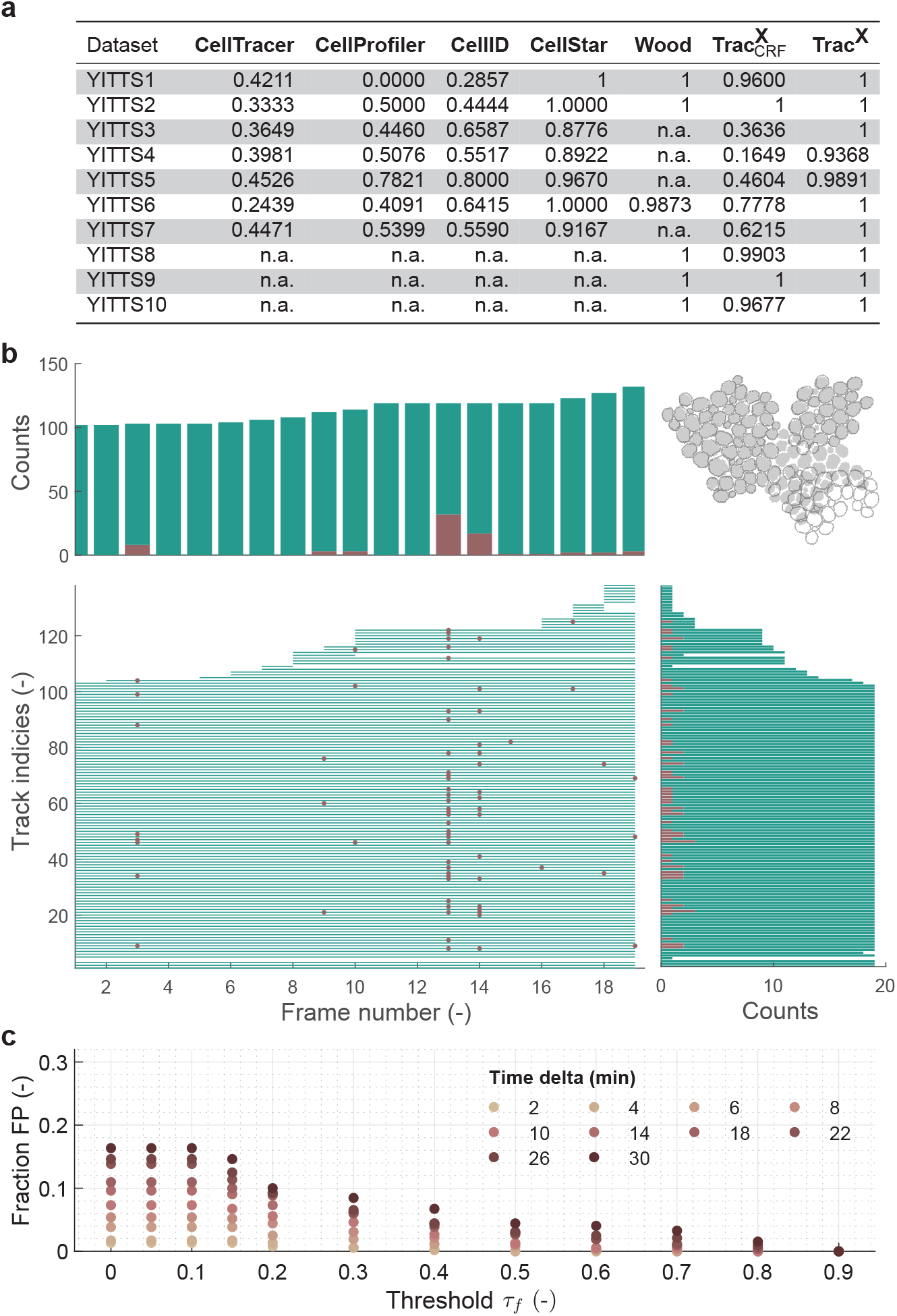
Performance of *Trac*^*X*^. **a** Performance assessment (long-term tracking quality characterized by the F-score) of *Trac*^*X*^ based on yeast image toolkit data sets YIT-TS1-10 compared to the five best-performing algorithms^10^ on these data. See **Methods** for details. Compared to the final implementation (*Trac*^*X*^), 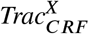 bases assignments only on the fingerprint distance (*d*_*f*_). **b** *CRF*-based track assignment score for each cell (track index) over time (image frames) with corresponding c^*f*^ounts (top and right histograms) in *S. cerevisiae* data set YIT-TS3. Green lines: *d*_*f*_ is lowest for the given assignment; red dots: cells with lower *d*_*f*_ exist, indicating uncertain assignments. Top right: example segmentation masks for the transition from frame 12 (outlines) to frame 13 (filled). **c** Fraction of false positives (FP, assignments identified as questionable despite being correct; here normalized by the total number of assignments) for yeast data set TS-SC9 as a function of neighborhood threshold *τ*_*f*_ and imaging frequency.

We also intended *CRF*-based classification to support targeted manual curation of results from any tracker—manual verification of each assignment is impractical for large data sets with thousands of linkages. *Trac*^*X*^ calculates post-tracking *F*_*f*_ values for all assignments to highlight potentially problematic cell tracks or image frames. For the perfectly tracked (compared to the manual ground truth) data set YIT-TS3, classification with *τ*_*f*_ = 0 indicates 72 out of 2286 assignments as problematic (**Fig. 4b**). For example, it highlights potential issues in frames 12-14, which are challenging for tracking because two colonies merge and move. These are false positives, but their number is comparatively low. We tested this systematically for a yeast data set with many cells and high imaging frequency (TS-SC9; see **Table S2**). **Fig. 4c** shows fractions of false positives (and corresponding efforts for manual curation) in low percentages for conservative threshold choices. In addition, it emphasizes the importance of a sufficiently high imaging frequency, adapted to the tracking problem (see also **Fig. S4**). Hence, *CRF*s help for high-precision automatic cell tracking and for efficient manual curation.

### Genealogies can be reconstructed with minimal additional information

To study cellular processes and their inheritance over multiple generations, one needs to reconstruct the genealogy (or lineage). It is represented as a tree where the tracks of the offspring are linked to the tracks of their mothers. Since we obtain tracks of high quality, we reasoned that we can create heritage assignments with minimal additional input. For this purpose, we developed algorithms for different cell types and cell division types (see **Methods** for details).

For asymmetrically dividing cells such as *S. cerevisiae*, we require an additional fluorescent marker that links mother and daughter cells. Here, we used Myo1, a protein localizing to the bud neck^31^. We determine assignments from the overlap of the marker with potential mother and daughter cells on each frame (**Fig. 5a**) and verify them using line profiles between mother and (future) daughter centroids (**Fig. 5b,c** and **Fig. S3**). With our own *S. cerevisiae* data (SC-TS7; see **Methods**), the algorithm correctly assigned all mother-daughter pairs (two seeding cells divided 122 times to a final 245 cells **Fig. 5d** and **Movie S1**).

**Figure 5.**
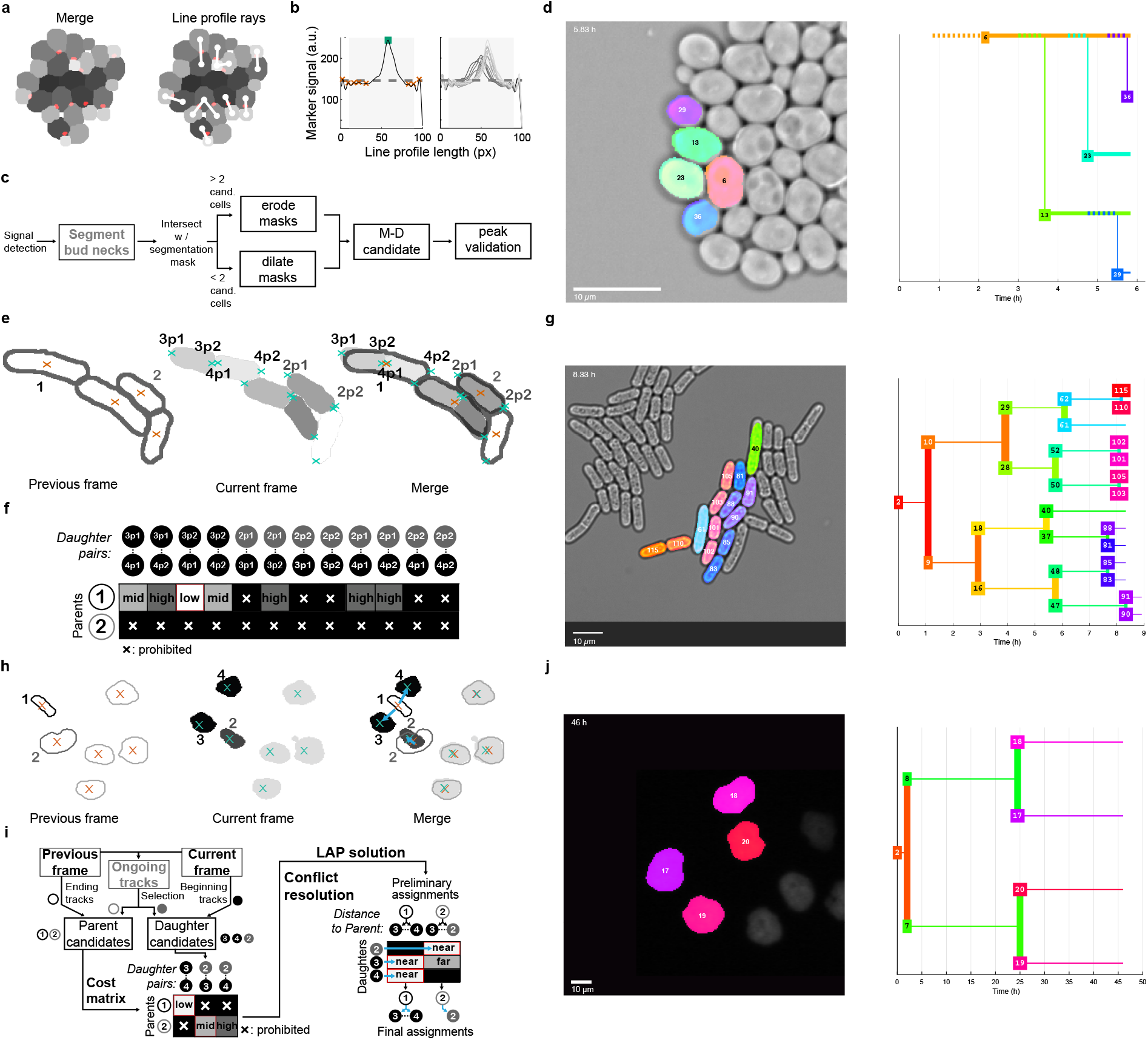
Genealogy reconstruction. **a-d** Lineage reconstruction for asymmetrically dividing *S. cerevisiae* cells using the bud neck marker Myo1. **a** Left: merge of segmentation masks for cells (grey) and marker (red). Right: overlay of profile lines (white) between centroids (white dots) of candidate mother-daughter pairs. **b** Left: typical, re-scaled marker signal between two cell centroids along the profile line (black line) with detected peaks (red crosses) and selected peak indicating bud neck marker presence (green square). Boundary regions (no gray background) are ignored. Right: marker signals for all cells in **a. c** Flowchart of the reconstruction. For each marker signal, neighbouring cell masks are eroded or dilated up to a single candidate mother-daughter cell pair, which is then validated by marker presence. **d** Cell lineage for a selected mother cell and all its offspring; cell identities (numbers, colors) match between example image and lineage tree. **e-g** Lineage reconstruction for symmetrically dividing, rod-shaped cells such as *S. pombe* using cell centroids and poles. **e** Segmented cells (black outlines and indices) and their centroids (red crosses) of the previous frame are merged with segmentation results of the current frame, including identified poles (cyan crosses; labels ‘*x*p*y*’ specify cell *x* and pole age *y*). **f** Schematic cost matrix for the daughter pair - parent assignment based on the distance between pole coordinates of cells in new tracks and centroids of cells in the previous frame. **g** Example lineage tree as in **d. h-i** Lineage reconstruction for convex, symmetrically dividing cell types such as N2DL-HeLa cells^33^ based on (nuclear) fluorescent label. **h** Similar to **e**, but with consideration of centroids in previous (red crosses) and current (blue crosses) frame. Black: candidate daughter cells. **i** Flowchart for reconstruction, example cost matrix, and assignment results after conflict and linear assignment problem (LAP) solution. **i** Example lineage tree as in **d**. Scale bars: 10 *μ*m.

For rod-shaped or other sturdy, symmetrically dividing cells such as fission yeast or bacteria, the algorithm tries to solve the genealogy using simple geometry, without additional input. Assignments are based on the shortest distance between the poles of the potential pair of daughter cells and the centroid of the mother cell (**Fig. 5e,f**). Thereby, the genealogy and the pole age can be determined directly. We tested the algorithm using *S. pombe* data (**Fig. 5g**) as well as a published ground truth data set of growing *Bacillus megaterium* cells^32^ (see details in **Table S2**). We obtained error-free lineages (4 seeding cells divided 66 times to a final number of 111 cells. All 111 tracks were correct (**Fig. 5g, Movie S2**), and 8 seeding cells divided 92 times to a final number of 160 cells; all 160 tracks were correct (**Movie S3**).

For amorphous or convex-shaped, symmetrically dividing cells such as HeLa or embryonic stem cells, we modified this algorithm because one cannot easily determine cell poles. Without conserved cell shapes, we assume area conservation to solve the lineage via a linear assignment problem that incorporates distances between centroid positions as well as differences in cell areas into the cost matrix (**Fig. 5h,i**; see **Methods** for details). For a subset of test data set of mammalian (HeLa) cells^33^, we again obtained the correct lineage (4 seeding cells, 8 divisions, 20 final cells with correct tracks, 5j, **Movie S4**; recall for the full data set of 0.89 (236 / 264 cells correct), **Movie S5**). Overall, this supports versatility and precision of *Trac*^*X*^ also for lineage reconstruction.

### Lineage analysis suggests a comprehensive concept for cell size homeostasis in *S. cerevisiae*

To demonstrate the application potential of *Trac*^*X*^, we revisited the fundamental question how yeast cells control their size depending on nutrient availability. It has the aspect of how individual cells define the set-point for size control, specifically for leaving the G_1_ phase of the cell cycle (in a transition termed Start in *S. cerevisiae*) when they reach a critical size. Such size control with stochastic elements is well-established at the single-cell level^34^. An intriguing molecular hypothesis is that differential scaling of Start inhibitor (with Whi5 as a dominant actor) and activator (such as G_1_ cyclin Cln3) concentrations with cell volume establishes size control. In such models, high inhibitor concentrations allow new-born cells to grow sufficiently in G_1_ until the balance shifts to activators at Start^35–37^. Whi5 levels may also provide a memory of environmental conditions (growth rates), and thus modulate size set-points according to nutrient availability^38^. However, the other, largely elusive aspect is cell size homeostasis: irrespective of the presence of most known regulators, cell size distributions of yeast populations are equally narrow^37^. Because cell size control and homeostasis are highly multi-factorial^39^, focusing on (multiple) molecular regulators^35,36,40^, nutrient conditions^38,41–43^, or lineage analysis^44^ alone may therefore be too limiting.

Aiming for a more integrative analysis, we acquired single-cell and corresponding lineage data for *S. cerevisiae* strains carrying fluorescent markers for Whi5 and Myo1 and growing in three different glucose concentrations (0.05 mM, 0.2 mM, and 100 mM) in a controlled microfluidic setup for up to six generations (see **Methods** for details). **Fig. 6a** shows the long-term dynamics of an example yeast cell at the highest glucose concentration. Parameters obtained by image analysis include cell and bud (i.e., future daughter) volumes as well as spatially resolved maker concentrations. We defined the G_1_ phase as the time interval with nuclear Whi5, as in previous studies^34,37^. Because our interest was size control in G_1_, we subsumed the rest of the cell cycle under G_2_/M, noting that Myo1 marker and bud presence allow for a finer discrimination in principle. Corresponding cell cycle annotations as well as the inference of cell states and rates were automated (see **Methods** and **Fig. S11a** for details).

**Figure 6.**
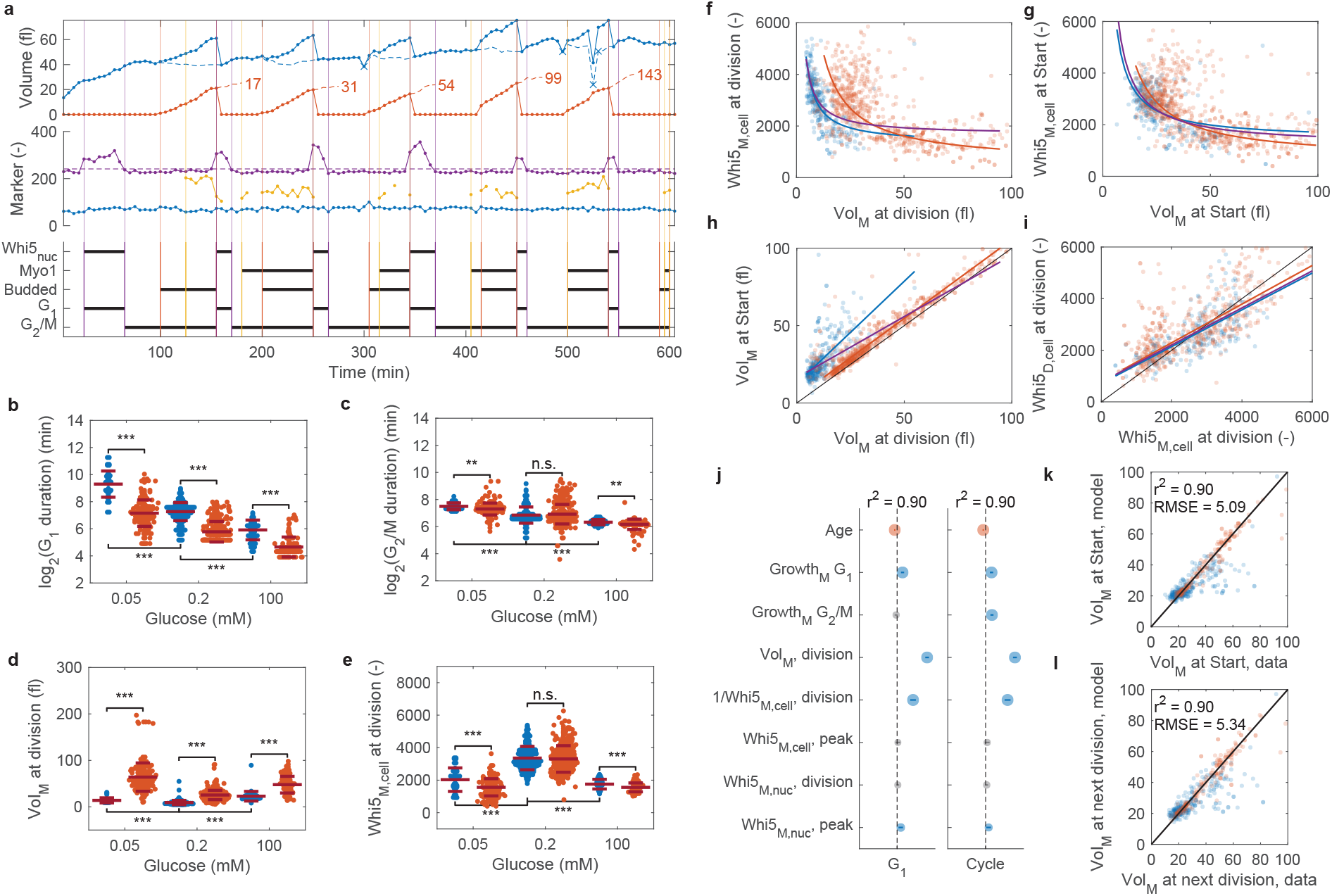
Effects of glucose availability on cell cycle regulation of *S. cerevisiae*. **a** Example single-cell track for growth on 100 mM glucose. Volumes: mother including (blue solid) and excluding (blue dashed) bud, bud (red solid), and initial volume dynamics of daughter cells (numbered) after division (red dashed). Markers (arbitrary scaling): Whi5 nuclear concentration normalized by Whi5 cellular concentration (purple), Whi5 cellular concentration (blue) and bud neck marker Myo1 (yellow). Automatically identified phases are indicated by black horizontal lines in the lower panel, and corresponding vertical lines in all panels. Subscript: nuc, nuclear concentration. **b-e** Cell cycle states grouped by newborn (blue) and older (red) cells and glucose concentrations. Purple lines show mean ± s.d. Results of unpaired two-sided t-tests: *, *P* < 0.05; **, *P* < 0.01; ***, *P* < 0.001; n.s., not significant (*P* > 0.05). From left to right, group sizes for complete cell cycles were *n* = 41/176/189/283/54/84, and for additional G_1_ phases *n* = 2/22/32/116/1/30. **f-i** Correlations of cell cycle variables for newborn (blue) and old (red) cells with data (points) and corresponding regressions (lines; inversely proportional for **f,g** and linear for **h,i**). Purple line: regression for all cells. Subscripts: M, mother; D, daughter; cell, cellular concentration. **j** Effect sizes in linear models for cell size at the end of G_1_ (volume at Start) and of the cell cycle (volume at next division). Symbols: significant negative (red), significant positive (blue), and not significant (grey) at *α* = 0.05. Sizes are proportional to −*logP* for two-sided t-tests of coefficients. All tested covariates are shown. Vertical bars show 95% confidence intervals of estimates. **k,l** Model predictions vs data for individual newborn (blue) and older (red) cells. Black line: identity. RMSE: root mean squared error.

Regarding cell cycle phases, daughter cells (newborn cells with replicative age 0) spent significantly longer time in G_1_ than mother cells (replicative age > 0) and G_1_ duration decreased consistently with increasing nutrient availability (**Fig. 6b**) and correspondingly increasing single-cell growth rates (**Fig. S11b**) as previously reported^41^. Also G_2_/M duration decreased significantly with nutrient availability, albeit not always between mothers and daughters (**Fig. 6c**). This matches earlier observations with growth modulation by different carbon sources^40,42,44^ and suggests that it is not simply the nutrient type that sets the G_2_/M duration.

Cell volumes at division showed a non-monotonic relation with nutrient availability: for both (future) mother cells at division (**Fig. 6d**) and their daughters (**Fig. S11d**), cells at the intermediate glucose concentration were smallest on average. Increased volumes at low growth rates were not observed in recent studies focused on Whi5^38,41,42^; we assume this is because they did not cover as long doubling times as our lowest glucose concentration (on average ≈16h and ≈6h for newborn and older cells, respectively). Our experiments, however, are consistent with a classical study’s results on minimal media^45^. This non-monotonic relation could explain at most weak linear associations between growth and cell size observed in yeast populations^46^ and it emphasizes the need for single-cell analysis along lineages.

Because total Whi5 levels are positively correlated with G_1_ duration in different nutrients in a mother- and daughter-specific manner^37,38,41^, we asked if a similar relation exists between cellular Whi5 concentration and cell volume at division. Consistent with previous data^41^, and except for the medium glucose concentration, these Whi5 levels were significantly higher in daughter than in mother cells (**Fig. 6e**). However, the most important difference existed between nutrient conditions, suggesting a negative association between cell volume and Whi5 levels that is not nutrient-specific (**Fig. 6d,e**, and similar for nuclear Whi5 in **Fig. S11d**). At the single-cell level, volumes at division and cellular Whi5 concentrations at division showed inverse relations that were clearly separated for mother and daughter cells (**Fig. 6f**). These two distributions, however, merged primarily via daughter cell growth when considered at Start (**Fig. 6g**), in contrast to the separation between mother and daughter cells when considering the volumes at division and Start (**Fig. 6h**). Together with the high percentage of variance explained by the Whi5-volume relations (*r*^2^ > 0.4, *P* < 10^−10^ in all cases), this indicates that cellular Whi5 could be an important determinant for the cell size set-point in G_1_.

To test this hypothesis, we estimated linear models to predict cell volumes at Start and at the end of the cycle, using the inverse of the cellular Whi5 concentration at division together with other plausible variables as predictors (see **Methods**). Both models provided excellent fits to the data (**Fig. 6j-l**). To predict the volume at Start, the model indicates independent and important effects of the volume and the Whi5 cellular concentration at the start of G_1_ (**Fig. 6j**). Growth in G_1_, cell age and nuclear Whi5 concentration have only minor effects, and factors such as growth in G_2_/M (a negative control) do not reach significance. We observe a similar pattern for the entire cycle, providing further evidence for the predictive value of Whi5 concentration for the volume set-point in individual cells.

With respect to cell size homeostasis and Whi5’s role in it, it was suggested that Whi5 levels memorize growth rates in earlier cycles, based on population data^38^. For single cells, we do not find such a correlation (**Fig. S11e**), and only weak positive correlations between volume at division and mother or bud growth rates (**Fig. S11f,g**). Memory of growth rates, in addition would not achieve size homeostasis because it would represent a positive feedback. Our data suggests that size homeostasis involves a return to the mean in a stochastic process (**Fig. 6i**); daughters of small mothers tend to have higher Whi5 concentrations than their mothers and thereby a higher volume set-point. The inverse holds for larger-than-average mothers, leading to a return to the mean size over successive generations. Clearly, we need additional factors to explain increasing daughter volumes with nutrient availability (**Fig. S11d**) and the duration of cell cycle phases is less well predicted than cell volumes (**Fig. S11i,j**) without incorporating activators such as cyclin Cln3^41^.

To estimate Whi5 concentrations, we followed previous studies^38,41^ by normalizing total cell fluorescence with cell area. Alternatively, one can normalize by cell volume^35^. This concentration estimation had only two effects (**Fig. S12**): reducing the impact of cellular Whi5 on cell size predictions (**Fig. S12g**), and not providing evidence for the hypothesis on size homeostasis in the population (**Fig. S12e**). Hence, quantification methods for protein (Whi5) concentrations require future, detailed investigations. Overall, however, our multi-dimensional analysis of single-cell behaviors purely in glucose media suggests that Whi5 concentration in the cell provides the volume set-point for yeast cells at Start, with Whi5 distribution along the lineage as a hypothesis for how size homeostasis of the population can be achieved.

## Discussion

High-quality segmentation and tracking is key for any analysis of microscopy data. However, tracking is difficult due to different imaging modalities, changing growth environments, and different cell types. This makes quality control critical, so far by tedious manual curation that does not scale. Cell region fingerprinting (*CRF*) overcomes this limitation because it reliably discriminates between correctly and incorrectly assigned cells on consecutive images without ground truth or fine-tuning of parameters. Importantly, we showed that *CRF*s generalize to different cell types and imaging modalities. Their major limitations are dealing with fast-changing regions (which can be well-controlled by adequate imaging frequency) and ‘unnecessary’ manual curation when false positives occur (but these are a few percent of all assignments and they do not influence the tracking performance). An avenue for future development is to extend the *CRF* to higher spatial dimensions.

Our tracker *Trac*^*X*^, which is based on established ideas but includes our new automated *CRF*-based proofreading concept, underlines both generality and performance of the concept. For example, it is not trivial that a ‘simple’ tracker out-competes established algorithms also for very challenging problems, while additionally flagging suspicious results at the single-cell level. Apart from general limitations due to inaccurate inputs (primarily bad segmentation results), specific limitations arise again from fast-changing neighbourhoods. However, the latter can be overcome by good experimental designs to keep the cells in position, for example, with microfluidic culturing devices or increased imaging frequency. Highly accurate tracking then also helps to reconstruct the genealogy with high precision for symmetrical and asymmetrically dividing cells. Importantly, the modular design of *Trac*^*X*^ makes it a valuable tool in image analysis pipelines, for example. Because one can choose the-best suited segmentation algorithm for each cell and division type. Alternatively, the *CRF* classifier can be applied with other tracking algorithms to subset or manually curate tracking data. One could, for example, thus efficiently create training sets for neural networks or increase the statistical power in biological applications.

Finally, *Trac*^*X*^’s capabilities enabled us to revisit the classic question how yeast cells control their size depending on nutrient availability. Its output at the single-cell (lineage) level highlights complex relationships between nutrient availability, replicative age, cell cycle progression, and cell size control, Specifically, the proposed model for Whi5-mediated control of cell size in single cells and at the population level may reconcile largely contradictory findings (or their interpretation based on few parameters) in previous studies. Evidently, future analyses are needed, for example, to assess in detail if dilution of Whi5 is central to size control^35^ or not^36,39^, and to what extent different methods of protein concentration estimation impact such conclusions. Such analyses will also need to include activators such as Cln3, the counterparts of Whi5 and related inhibitors^37^. Overall, we argue that our application to cell cycle control of *S. cerevisiae* shows the need for better, more accurate analysis methods for live-cell imaging, such as achieved with *Trac*^*X*^. We are confident that the possibility for mostly automated single-cell analysis along lineages at large scale will allow further biological insights well beyond similar, cell cycle related questions.

## Methods

### General notations

We define a time-lapse microscopy dataset as an ordered set of *T* images (or image frames) indexed by *t* = 1 … *T*. An image contains a set 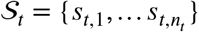 of segmented objects (in our case, cells, which we use interchangeably). Note that different images can have different numbers *n*_*t*_ of such objects. We denote by 𝒮 = ⋃_*t*_ 𝒮_*t*_ the set of all segmented objects over all images. Finally, we define the cell neighborhood of a segmented object *s*_*t,i*_ in the same image frame *t* as the set 𝒩 (*s*_*t,i*_) ⊂ 𝒮_*t*_ of objects adjacent to *s*_*t,i*_. Adjacent objects are those objects directly surrounding *s*_*t,i*_ in a fixed perimeter, which is a user-adjustable parameter.

### Object features

For each segmented object, various features such as the object centroid position, the object’s area covered and orientation in the image, and potentially quantified signals from additional channels (for example, fluorescence) are used. Internally we use the features defined by CellX^21^ as well as newly defined features for symmetrical lineage reconstruction (coordinates and ages of cell poles, cell orientation, and cell major axis). All segmentation masks from other algorithms are converted into this format prior to tracking.

### Cell region fingerprint and fingerprint distance

The cell region fingerprint (CRF) is calculated as the precise descriptor of the local image information around a segmented object’s centroid position on a given image. We apply a cell region fingerprint window with constant side length (*f*_*l*_; see **Table S1** for all methods parameters) to extract a squared subset image matrix from the raw image. It is resized by a constant factor (*f*_*r*_). The resulting matrix is Fourier transformed using the discrete cosine transformation (DCT). From the DCT transformed matrix, the coefficients for the *f*_*q*_ lowest frequencies (DCT coefficients) form the CRF denoted by *f* (*s*_*t,i*_), where we set the direct current (DC) coefficient at (0,0) to zero due to its high intensity compared to the remaining frequencies. Hence, *f* (*s*_*t,i*_) is a *f*_*q*_ ×*f*_*q*_ matrix.

We measure the similarity of two objects (and their associated regions) by the cell region fingerprint distance *d*_*f*_ ∶ 𝒮 ×𝒮 → ℝ. It is defined as the normalized Euclidean distance of the objects’ CRFs. For a pair of objects (*i, j*) in consecutive frames:

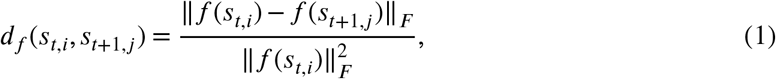

where ‖*M*‖_*F*_ denotes the Frobenius norm of matrix *M* with elements 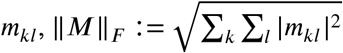.

### Neighborhood fraction

Our criterion for the classification of cell-to-track assignments is based on the fraction of objects in the neighborhood of *s*_*t*+1,*j*_ that have a lower cell region fingerprint distance to the assigned object in the previous frame, *s*_*t,i*_, than *s*_*t*+1,*j*_ itself. Formally, with the subset of neighborhood objects and of their subset with lower *d*_*f*_ specified as:

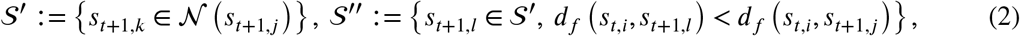

this neighborhood fraction is:

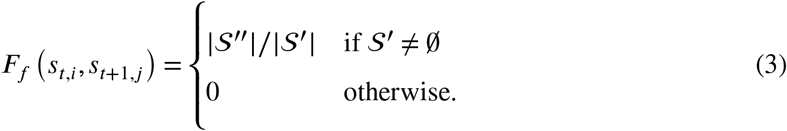

We classify assignments using a constant threshold *τ*_*f*_, where *F*_*f*_ ≥ *τ*_*f*_ indicates an incorrect assignment.

### Initial cell-to-track assignment

We obtain initial cell-to-track assignments by solving a linear assignment problem (LAP) with the Jonker-Volgenant algorithm that minimizes the total assignment cost^24^. For frame *t* > 1, assignments are made between two sets of objects: cells in the previous frame together with previously unassigned cells, denoted by 𝒮^*S*^ = 𝒮_*t*−1_ ∪ 𝒮^*O*^ with elements 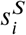, and cells in the current frame, denoted by 𝒮^*T*^ = 𝒮_*t*_ with elements 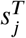. The set 𝒮^*O*^ is initialized to be empty and changes during iterations by adding unassigned cells and removing elements according to the number of frames a cell remained unassigned (see below). To account for frame skipping, we define *σ*(*s*) for a cell *s* as the difference between the current frame *t* and the last frame with assignment.

The elements of the |𝒮^*S*^| × |𝒮 |^*T*^ total cost matrix *C* result from contributions accounting for cell displacement, cell size deviations, cell rotations, and frame skipping. Because only subsets of the contributions may apply for an assignment problem, we use the general formulation:

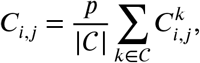

where 𝒞 is the set of partial cost functions. The normalization makes costs comparable with different numbers of active partial costs, and the proportionality constant *p* is adjusted for numerical efficiency in solving the LAP (typically, we adjust *p* to obtain comparable maximal partial cost function values).

To compute the cost for cell displacement, let the vector-valued function **x**(*s*) be the centroid positions of a cell considered for assignment (**Fig. S5a,b**). We predict movement of cell 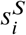 from the last preceding frames with assignment (e.g., *t* − 2 and *t* − 1), assuming that it is conserved; the vector function *δ*(*s*) denotes the estimated displacement filtered by the neighborhood 𝒩 (*s*). We use a simple Gaussian filter with a custom radius (default value 100 for × and y) to down-weigh displacements of distant neighbours. The displacement cost matrix is then defined as:

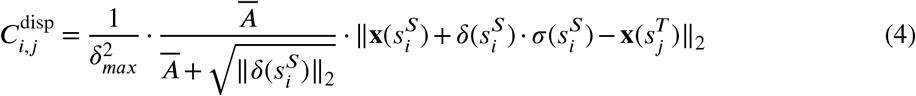

With the first term, centroid displacements are weighted symmetrically such that a unit cost is reached at the maximum allowed displacement *δ*_*max*_ (**Fig. S5B**). This parameter has a physical interpretation: it is the distance (in pixels) a cell centroid may move under normal growth between consecutive frames. This depends on cell type and acquisition system (e.g., magnification and pixel size). The second term relates the displacement to the average cell area, 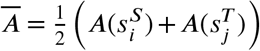, where *A*(*s*) gives the area of object *s*. The last term accounts for the distance between predicted and actual position of 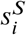 in frame *t*.

A second cost matrix captures changes in cell size, where cell size deviation is weighted asymmetrically to account for balanced growth (**Fig. S5c**). The underlying assumption is that a young cell increases its size rapidly, whereas loss in size is often due to mis-segmentation in a crowded region. However, we allow for enough flexibility to reflect situations where a cell’s size might change due to environmental perturbations such as salt stress. With the cell area ratio 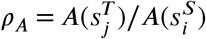 and the asymmetric expansion functions

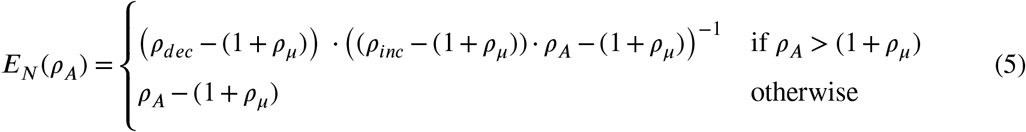

and

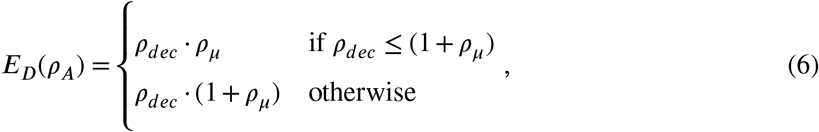

the cell size cost matrix is:

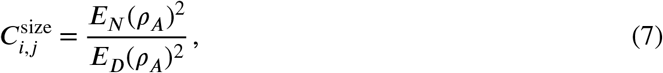

where the denominator adjust the costs around the growth rate. It uses parameters for maximal allowed relative size decrease (*ρ*_*dec*_) and increase (*ρ*_*inc*_) as well as an offset accounting for balanced growth (*ρ*_*μ*_; see **Fig. S5c**).

A cost matrix that takes cell rotation into account can help tracking cell types with non-convex shapes.

We denote by *θ*(*s*) the orientation of cell *s* (in degrees). With the rotation cost matrix

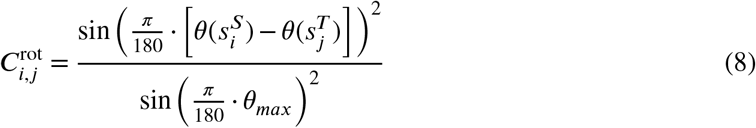

the rotation cost is symmetric and defined such that the maximum allowed rotation angle (*θ*_*max*_) of the cell’s major axis on consecutive frames is where the cost reaches unity (see **Fig. S5d**).

Finally, the cost matrix for frame skipping is defined as:

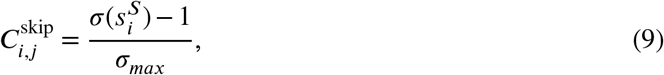

where *σ*_*max*_ is the maximum tolerated number of frames with a cell not being present (i.e., not segmented, see **Fig. S5e**).

As stated above, the final cost matrix is the sum of the individual weighted cost matrices and reflects the physical differences of cells between consecutive frames (see **Fig. S5f** for an example).

We also implemented a minimal version of the *Trac*^*X*^ termed 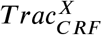. The aim of this tracker is to track only based on the *CRF*: the cost matrix uses only the cell region fingerprint distances between consecutive frames.

### Assignment validation and refinement

After solving the LAP using the above cost matrix, we get the best possible assignment for most of the cells. As a first independent proofreading step, we validate assignments by a neighborhood fraction *F*_*f*_ (Eq. 3) below the threshold *τ*_*f*_. Valid assignments are kept and invalid assignments are deleted for re-evaluation.

We refine assignments by first estimating the vector field of cell motions from validated assignments and then correcting or adding assignments iteratively by solving LAPs^26^ (see **Fig. S6**). During two refinement iterations, we first use Eq. 4 with its quadratic penalty, then we replace the displacement cost matrix by:

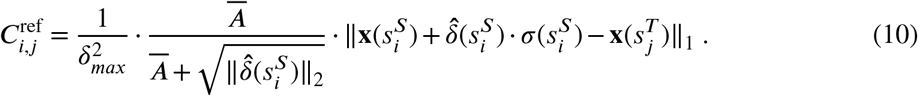

It allows for linear motion up to a defined distance (see **Table S1**) and uses the estimated vector field 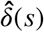. Next, we again use a modified Eq. 4 with a linear penalty to allow displacement only along the vectors.

Finally, we update the set of cells considered for future assignments with the set of unassigned cells up to frame *t*, 𝒮^*U*^, but remove cells that exceed the threshold *σ*_*max*_ for frame skipping, that is, 𝒮^*O*^ ← 𝒮^*U*^ (*σ*(*s*^*U*^) ≤ *σ*_*max*_).

### Lineage reconstruction

To determine the parent of each symmetrically dividing cell with rod-shaped phenotype, we calculate the cell pole coordinates *Po*_1,2_ from the cell’s centroid *s*_*x,y*_, major axis length *l*_*ma*_ and orientation *θ* via:

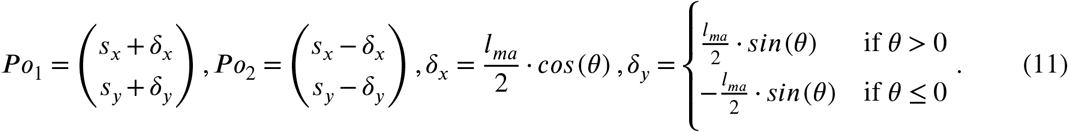

We then minimize the Euclidean distance between poles of newly starting tracks and the cell centroid of tracks ending in the previous frame within reasonable distance (defined by the parent cell length) and overlap with the parent track. This directly establishes the parent relationship for newborn tracks as well as the cell pole ages (poles with minimal distance are set to zero, their opposites increase age by one). Cell pole ages in the first frame are undefined.

To determine the parent of each track for symmetrically dividing cells with amorphous or convex phenotype, we assume that a cell division of one ending parent track leads to the beginning of two new daughter tracks. Because this might not always apply (e.g., if a track is a continuation of a parent track with one daughter cell) *Trac*^*X*^ tries to detect cases with strong size decrease. Possible daughter candidates are evaluated iteratively, while infeasible ones (spatial distance above a threshold or both daughters in a continuous track) are excluded on consecutive frames with an infinite cost. Assignments are made by solving the linear assignment problem with the LAPV. The solution can contain the same daughter in more than one daughter-parent assignment. To resolve that, each daughter is assigned to only its closest assigned parent. However this might break daughter pairs, now lacking a daughter. Finally, the daughter parent assignments are set by setting the parent for each daughter. Wrong continuous tracks are now marked for splitting and single daughter parent pairs are marked for joining. See flowchart in **Fig. 5i** for details.

To automate the bud (daughter) to mother assignment for asymmetrically dividing cells with a fluorescent bud neck marker, we segment the marker by first de-noising the raw images with a total variation regularization filter (termed after the authors, ROF)^47^. Remaining speckles are removed by median filtering with a 3×3 kernel. We then compute the differential of Gaussian (DoG) with a rotationally symmetric Gaussian lowpass filter of equal size (21px) with standard deviations *σ* = [1, 2]. The resulting image is filtered by a canny edge filter with a user-definable parameter (edge sensitivity threshold; see **Fig. S3**) to detect the contours of the rod-shaped bud neck signal. We fill the holes in contours and dilate the image with a disk of 1px size to remove pixel fragment noise, followed by another round of morphological dilation and erosion. We then mask with the cell segmentation mask to set the signal outside single cells and colonies to zero. The resulting image is converted into a labeled mask. We keep bud necks within a specified area range and track them with the same algorithm as the cells, but with independently adjustable parameters. Daughter-parent assignments are performed iteratively on consecutive frames by optimizing bud neck overlaps with exactly two cell masks until convergence. The corresponding two tracks form a daughter-parent candidate if we also detect the bud neck signal independently on a line profile between the centroids of daughter and parent. Finally, we compute the most likely daughter parent pair as the one with the highest occurrence probability throughout its detection in experiment.

### Implementation

We implemented *Trac*^*X*^ in Matlab (MathWorks, Natick, MA) with a modular design, a command line interface (CLI), and a simple graphical user interface (GUI). Key features are: (i) *Trac*^*X*^ requires segmentation results in the format established by CellX; a module provides format conversions. (ii) One can test and modify default parameters on a subset of the data before batch processing. (iii) *Trac*^*X*^ provides tools to detect, visualize, and exclude segmentation artefacts such as cell debris and constrain input data by defining regions of interest, image border offsets, and thresholds on any segmentation feature, including fluorescence. (v) Tracking results are saved in tabular form, as control images for each frame to inspect the results, and as *Trac*^*X*^ state to resume work later. (vi) Image processing and visualization modules enable display and export of, for example, animated movies and genealogy. (vi) The modular design allows to extend *Trac*^*X*^ as well as its use in larger workflows (*Trac*^*X*^ saves project details as machine-readable XML files). For details, see **Supplementary Text**, methods parameters in **Table S1**, algorithm descriptions in **Figs. S7–S8**, and the user documentation at https://tracx.readthedocs.io/en/latest/.

### Performance evaluation

Performance evaluations focused on our *CRF* method’s ability to detect incorrect assignments in tracking. We therefore classify assignments between objects in consecutive frames as positive (predicted incorrect) if *F*_*f*_ > *τ*_*f*_, and negative (predicted correct) otherwise. To measure accuracy of the binary classification with respect to the ground truth (manual tracking), we use the balanced F-score (*F*_1_), the harmonic mean of precision and recall:

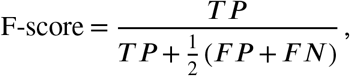

with true positives (*T P*), false positives (*F P*), and false negatives (*F N*).

To evaluate parameter influences on classifier performance, we determined F-scores for all combinations of window side length *f*_*l*_ ∈ [20, 30, 40, 50, 60, 80, 100, 120, 150, 200, 300] (px), re-size factor *f*_*r*_ ∈ [20, 25, 32, 40, 50, 64, 80, 121, 150, 200, 300] (px), and number of DCT coefficients *f*_*q*_ ∈ [2, 4, 8, 10, 12, 16, 20, 25, 30, 35, 40] (-). We used the following imaging data sets (see **Table S2**): 60mrnaCropped^48^, TS-SC9 (brightfield and fluorescence), and Fluo-N2DH-GOWT1^33^.

To assess *Trac*^*X*^ performance, we used the yeast imaging tool-kit (YIT)^11^ datasets summarized in **Table S2**. These ten datasets pose various challenges for cell tracking such as small colonies (YIT-TS1, 2, 6), colony translations and merging (YIT-TS3), and large colonies growing out of the field of view (YIT-TS4, 5, 7)^11^. We included three recent transmission light and phase contrast data sets that contain dense colonies and cells with irregular morphology (YIT-TS7-10)^11^,^30^. To obtain the missing segmentation masks, we re-segmented using CellX^22^, CellStar^11^ or YeaZ^49^ and then benchmarked *Trac*^*X*^ with the manually curated ground truth from YIT and their published evaluation platform (EP).

### Strains and cultivation

We used *S. cerevisiae* strains FRY2795 (a prototrophic haploid derivative of FY4 of genotype MAT*α* bearing Myo1-mKate (3x) and Whi5-mKO*κ* (1x) modifications inserted using an unpublished CRISPR protocol) and FRY2031 (an unmodified haploid prototrophic strain, FY4 of genotype MAT*α*). Details are given in **Tables S3-S5**. Cells were cultured in synthetic minimal media (SDmin) unless stated otherwise. We used 1.7 g l^−1^ yeast nitrogen base (BD Biosciences, Germany), without amino acids or ammonium sulfate, 5 g l^−1^ ammonium sulfate (Sigma-Aldrich Co., Germany), and 20 g l^−1^ D-glucose (Sigma-Aldrich Co., Germany) for the 100mM (2%) D-glucose condition. For the other media conditions with 0.2 mM (0.004%) and 0.05 mM (0.001%) D-Glucose, the liquid D-glucose stock was serially diluted for higher precision. Prior to imaging, cells were streaked on YPD plates at room temperature directly from the freezer. Single colonies were picked and diluted into the respective liquid media at 30°C in an orbital shaker days before the actual experiment. Using a Z2 Coulter Counter (Beckman Coulter, Nyon, Switzerland) and custom software, the liquid cultures were diluted to a cell density of 3 − 5 ⋅ 10^6^ cells per ml before loading them onto the microfluidic device (CellClamper^50^).

### Imaging and image analysis

In all experiments, we loaded the first two chambers with FRY2795 and the last two chambers with FRY2031, serving as growth control and to estimate background fluorescence. 3*μ*l of cell suspension were loaded into each chamber of the microfluidic device before covering it with a glass slide and connecting it to the media perfusion pumps. Pads of various sizes with at least one seeding cell were selected for imaging. During imaging, the respective medium was perfused with a flow rate of 15*μ*l/min by a Nemesys pump (Cetoni GmbH, Germany). A custom climate chamber around the Nikon Ti-E inverted microscope (Nikon Europe B.V., Amsterdam, Egg, Switzerland) kept the environment at constant 30° C. Medium reservoir and pumps inside the chamber kept the media at the same temperature. We equipped the microscope with a Nikon CFI Super Fluor 40X Oil, 1.3NA objective (Nikon Europe B.V., Amsterdam, Egg, Switzerland) and SpectraX light engine (Lumencore Inc., USA) light source. To excite the fluorophores, we assembled the following filter cubes: #1 EX: 600/14nm, BS: STHC 624nm, EM 655/40nm for mKate2 and #2 Ex: 546/6nm BS: ST565nm EM: 577/25nm for mKO*κ* (AHF Analysetechnik, Tuebingen, Germany). The imaging interval was chosen such that cells do not suffer photomorbidity^51^ based on the expected average cell cycle time (5, 12, and 10 min for 100, 0.2 and 005mM D-glucose). Microscope and peripherals were controlled by YouScope^52^. We performed at least two independent experiments per glucose concentration.

We used CellX^22^ for signal quantification and cell segmentation. For each experiment, a matching parameter file was tested on a few images to ensure best segmentation before batch analysis jobs were submitted using a custom job scheduler to a multi core machine (Power Edge T20 with Intel Xenon 2.1Ghz, 96 cores, 200 GB Ram, Dell Inc, USA). Segmented cells were tracked and assigned to a lineage using *Trac*^*X*^. Further analysis was performed with custom scripts in MATLAB (Mathworks Inc, USA).

### Cell cycle analysis

Per experimental condition (glucose concentration), we corrected CellX output for background fluorescence first by pixel-wise subtraction of the average fluorescence outside segmented cells and then by volume correction based on the control strain’s fluorescence. For the latter, we used a polynomial interpolant over all control cells. Concentrations of Whi5 in the respective cellular compartment (entire cell or nucleus) were then calculated as the total corrected fluorescence divided either by the compartment’s area or by its estimated volume. Volume growth rates were estimated for G_1_ and G_2_/M separately, using linear models (exponential models yielded the same estimation quality with our data).

To identify the phase of nuclear Whi5 (defined as G_1_ here), we derived a Whi5 signal by normalizing the median nuclear by the median cellular fluorescence to improve the signal-to-noise ratio. Then, per chamber, we fitted a two-component Gaussian mixture model to the Whi5 signal distribution and identified a common maximum likelihood threshold for classification of all corresponding cells and time points (see (**Fig. S11a** for an example). For the identification of budding and cell division events, we used the segmentation and Myo1 data as described above, except when Myo1 at the bud neck could not be detected (in that case, the start of Whi5 nuclear localization was used). Volume outliers were detected via the 1% quantile of cell volume differences between time points and discarded. Finally, we retained only complete cycles (showing the sequence cell division, G_1_, Start, budding, and subsequent division) and complete G_1_ phases (showing the sequence up to Start) for further analysis.

To test for significant differences between groups, unpaired two-sided t-tests for means were used without correction for multiple testing; we considered comparisons between conditions only significant when all individual group tests were significant at *α* = 0.05. For one-dimensional regressions, linear (*y* = *b*_0_ + *b*_1_ ⋅ *x* + *ε*) and inverse linear (*y* = *b*_0_ + *b*_1_ ⋅ *x*^−1^ + *ε*) models were used. For multidimensional models, we used robust linear regression (Matlab function `fitlm`) after discretizing the cell age into newborn and old (replicative age = 0 and > 0, respectively). For predictor variable selection, phase durations were not included because they directly depend on growth rates and volumes when a size set-point is assumed. For all regressions, unadjusted coefficients of determination (*r*^2^) are reported.

## Acknowledgements

We thank Markus Dürr for initial implementation of Ricicova’ tracker in R and Urs Küchler for its translation to Matlab. We thank Gregor Schmidt for training of the CellClamper and sharing datasets.

## Author contributions

**A.P.C**., **F.R**. and **J.S**. conceptualized the project. **A.P.C**. developed the CRF and wrote the TracX software. **A.P**. wrote the Gaussian vector filtering. **T.K** initiated and co-developed the graphical user interface and wrote the amorphous cell lineage reconstruction. **A.P.C**. engineered the cell strains and performed the microscopy experiments. **A.P.C**. and **J.S**. validated the software and analysed the data.**A.P.C**. and **J.S**. wrote the manuscript. All authors read and approved the final manuscript.

## Data availability

The data used in this study as well the analysis scripts are available for download at https://polybox.ethz.ch/index.php/s/iwOBUsjNFYIAqaX.

The *Trac*^*X*^ software can be downloaded from https://gitlab.com/csb.ethz/tracx as well as demo data from https://gitlab.com/csb.ethz/tracx_demo_data.

