## Supplementary Information for "Cell region fingerprints enable highly precise single-cell tracking and lineage reconstruction"

### Supplementary Figures

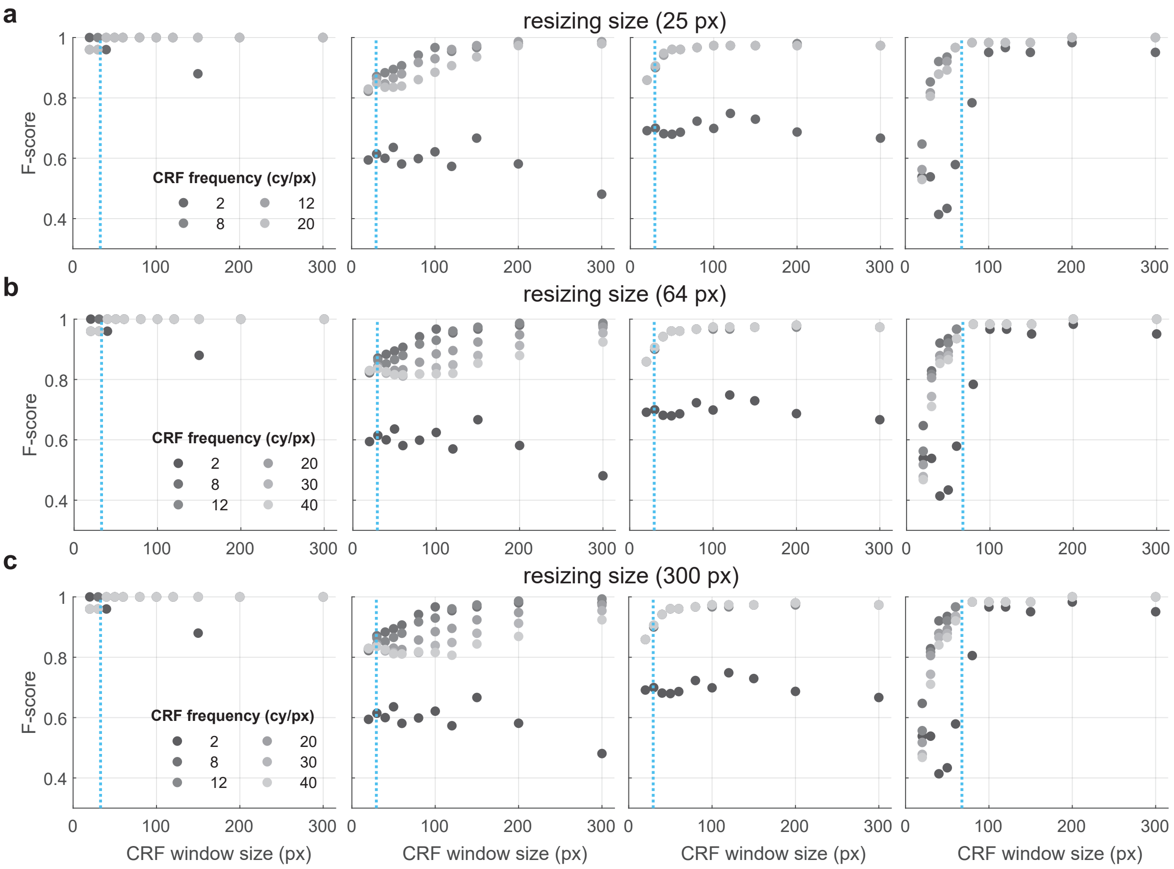


**Figure S1. The effect of the re-size factor on *CRF*-based classification.** $f_r$ values are indicated per row and data presented as in **Fig. 3b**.

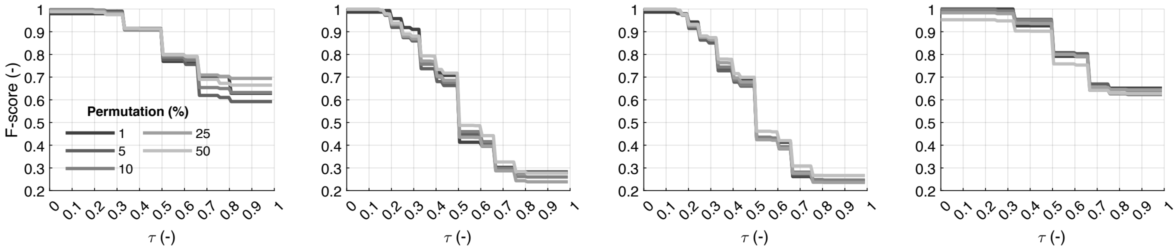


**Figure S2. Evaluation of the effect of the neighborhood fraction threshold ($\tau_f$) for the different cell types and image modalities in Fig. 3.** For each data set, we chose a random parameter set yielding a high F-score according to the data in **Fig. 3**. Window sizes ($f_l$), frequencies ($f_q$) and resizing ($f_r$) were [30, 8, 32], [300, 12, 40]. [300, 10, 25], and [200, 20, 32]. We permuted assignments by the fractions indicated. The very conservative value $\tau_f = 0$ can be relaxed to ≈0.2 while still accurately detecting matching consecutive (cellular) regions on all images and for all permutation frequencies.

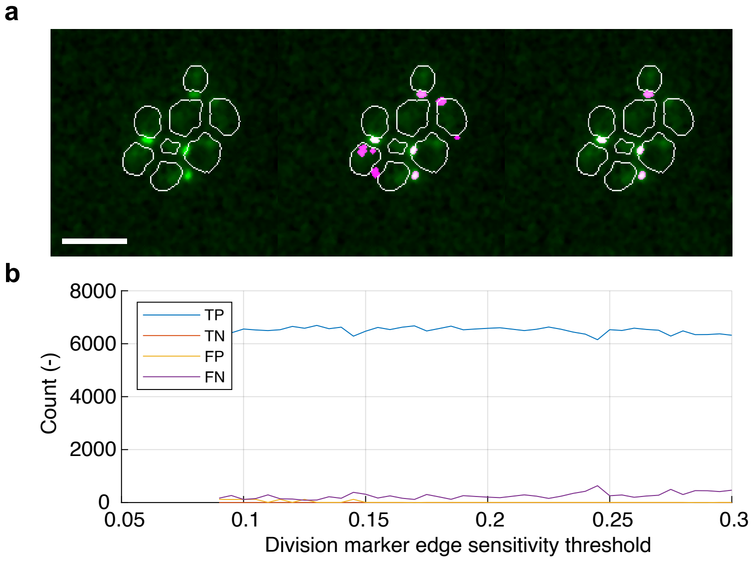


**Figure S3. The effect of the edge sensitivity threshold on asymmetric lineage reconstruction performance.** **a** Left: total variation regularization (ROF[1]) filtered false colored fluorescence image channel depicting the bud neck (green) with cell segmentation outlines (white). Middle and right: bud necks (magenta) detected for edge sensitivity thresholds of 0.09 (middle) and 0.225 (right). Scale bar = $10\mu m$. **b** Assignment counts as a function of edge sensitivity threshold classified into true positive (TP, bud to mother assignment correct), true negative (TN, no mother for bud), false positive (FP, wrong bud to mother assignment), and false negative (FN, missing bud to mother assignment).

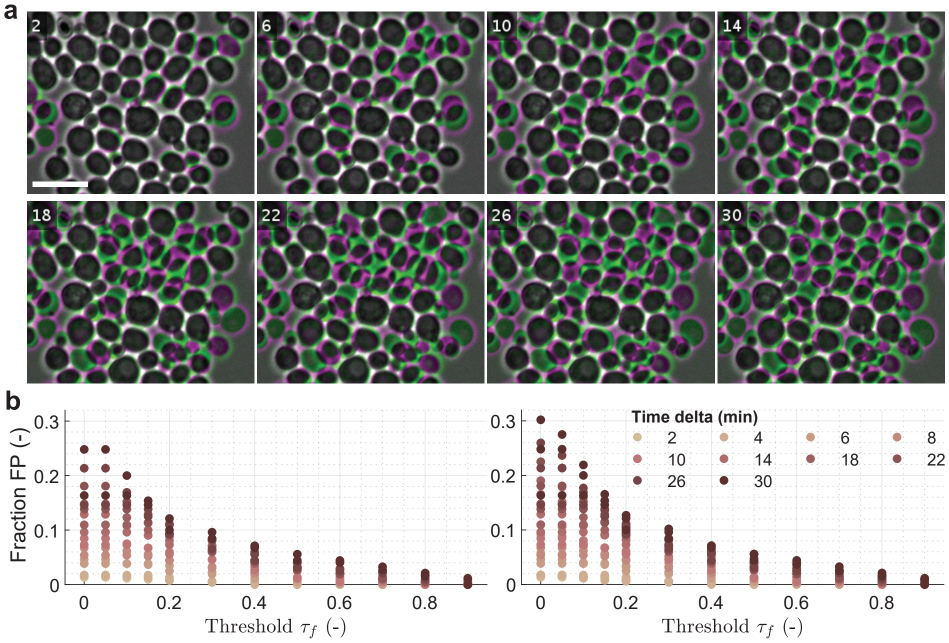


**Figure S4. Effect of imaging frequency on the false positive rate**. The relative rate of false positives ($F_f$) as a function of neighborhood fraction thresholds ($\tau_f$) evaluated for different imaging frequencies (colored dots). **a** False color image overlays of the first image frame with the one after 2, 6, 10, 14, 18, 22, 26, 30 minutes to simulate different imaging frequencies. Data-set used was TTS SC9 (see **Table S2**). Scale bar = $10\mu m$ **b** Fraction of false positive for the $F_f$ as a function of neighborhood fraction threshold ($\tau_f$). Neighbourhood window size was 60 px or 8.80 ± 0.14 neighbours (left panel). Neighbourhood window size was 80 px or 13.70 ± 0.31 neighbours (right panel). n=520 assignments.

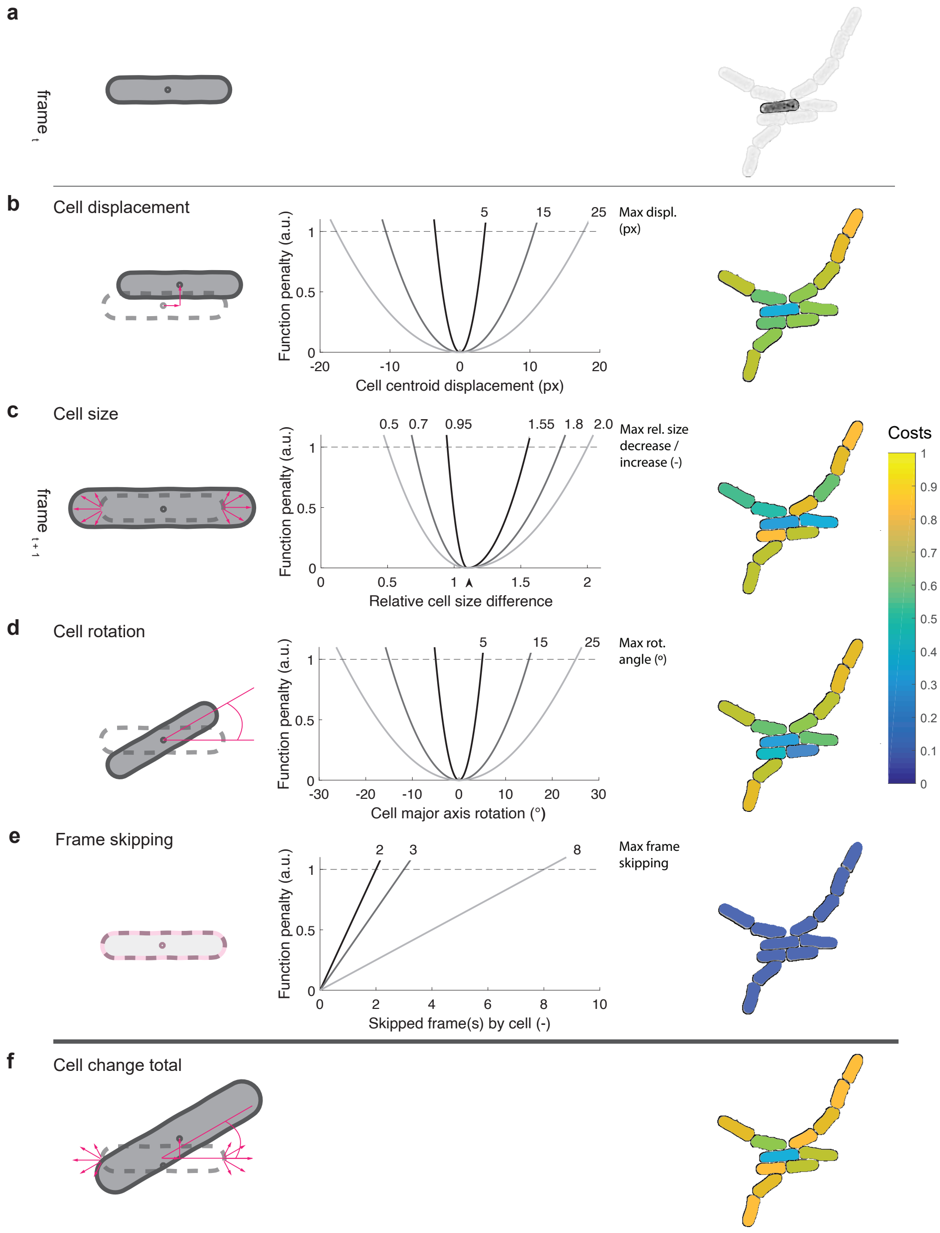


**Figure S5. Tracker cost matrix functions.** **a** Left: schematic cell in frame $t$. Right: real cell for which the costs will be computed (dark) and its neighbors (light). **b** Left: schematic cell in frame $t$ (dashed line) and at $t+1$ (solid) with cell centers (dark grey dots) and their translation (pink). Middle: relative displacement versus normalized cost. Solid lines correspond to the maximum allowed cell center displacement (x and y, in px) to reach unit cost (dashed line). Right: displacement costs for the cell and its neighbors (colors). **c** Left: as **b**, but with red lines indicating size differences. Middle: relative size difference versus normalized cost. Solid lines show maximum allowed cell size differences (decrease and increase, independently defined, in %) to reach unit cost, the arrow the offset to correct for average cell growth. Right: color-coded size costs. **d** Left: schematic cell with highlighted rotation of the major axis (red). Middle: difference in rotation versus normalized cost. Lines correspond to the maximum allowed major axis rotations (in degrees) for unit costs. Right: color-coded rotation costs. **e** Left: schematic cell with highlighted skipped frame (red). Middle: Frame skipping versus normalized cost. None of the cells skipped a frame (good segmentation), therefore all cells have cost of 0. **f** Left: total changes between frames. Right: color-coded sum of all cost functions. The highlighted cell from frame $t$ has the lowest total cost on frame $t+1$, which does not necessarily apply for the individual costs.

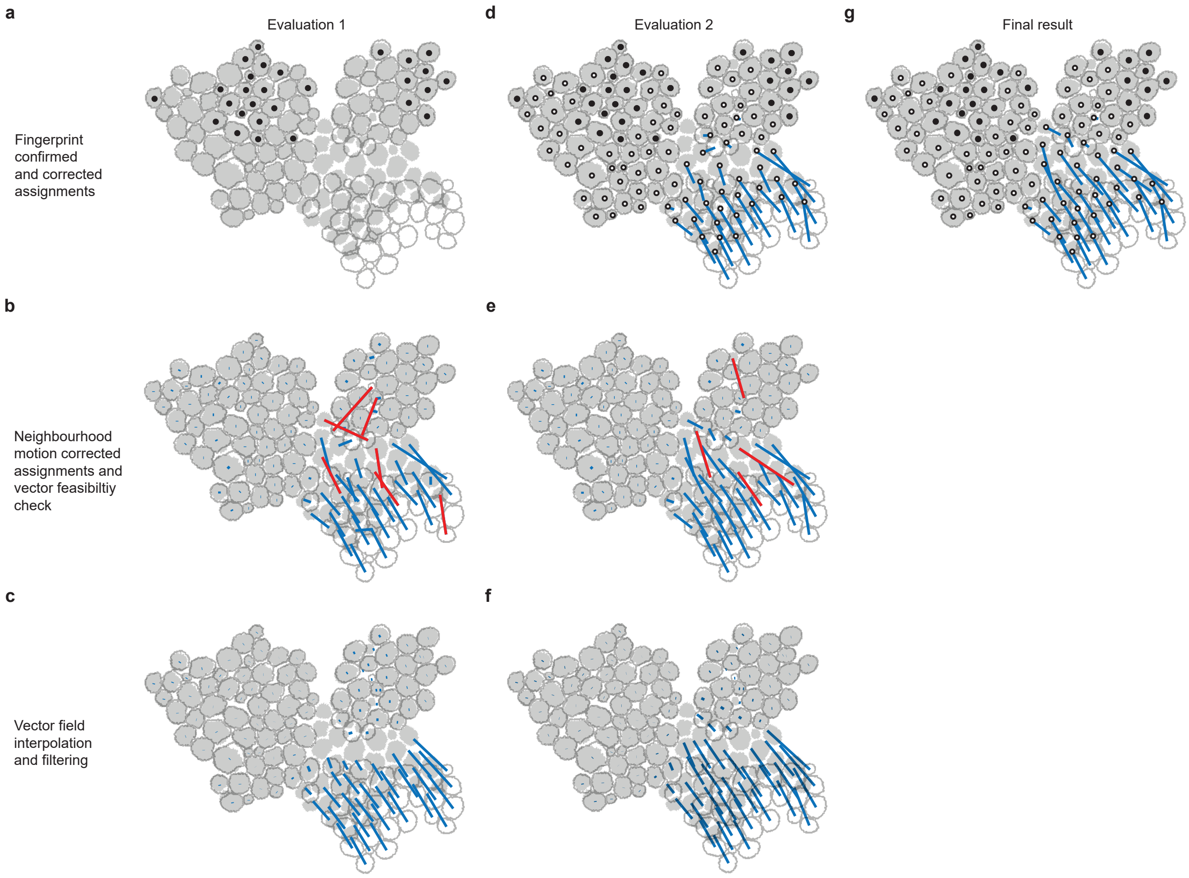


**Figure S6. Vectors of cell-to-track assignments during motion.** First (**a-c**) and second (**d-f**) evaluation step of the motion validation module. **a** Segmentation masks from $t-1$ (gray outlines) and from $t$ (filled gray) with assignments passing the proofreading step; cell centers marked as black dots. **b** Motion vectors of the assignments after solving the LAP that passed (blue) and did not pass (red) the feasibility check. **c** Motion vector after filtering out the unfeasible vectors and interpolating their motion based on the neighborhood (blue). **d** New assignments included in the second evaluation; cell centers marked as black circles. **e** Motion vectors after LAP solution as in (**b**). **f** Motion vectors after filtering and interpolation prior to the final assignment. **g** Final result showing the motion vectors (blue) for all assigned cells.

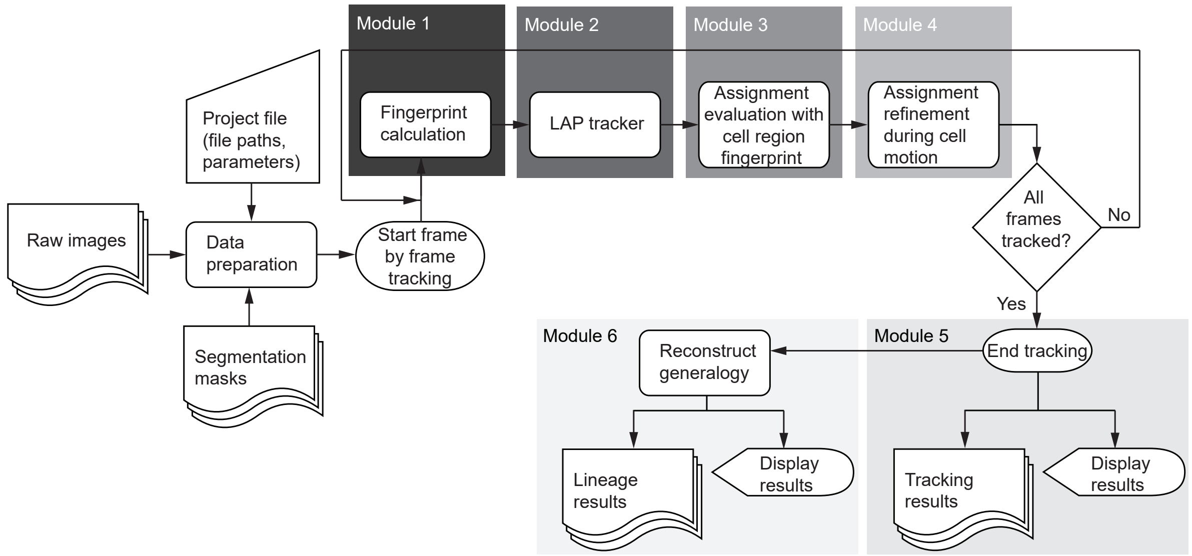


**Figure S7. Flow chart of the *Trac$^X$* core software architecture.** A tracking project defines all the required parameters and links to input files such as raw images and segmentation masks. Tracking is then started in a frame-by-frame manner. First, the *CRF* is calculated for each segmented centroid on each image frame. A first assignment is obtained from the LAP tracker process. These assignments are evaluated for correctness using the fingerprint distance ($d_f$), before refinements to handle unexpected motion where the image frequency was too low for the $d_f$ to be informative enough. When all frames were tracked, results are saved and displayed to the user. Boxes encode for manual user input (trapezoid), documents (waved rectangle), process (rectangle), decision (square standing on tip), start and end of the flow (ellipse), data display (ellipse with tip to the right), complying with the commonly used ISO 5807:1985.

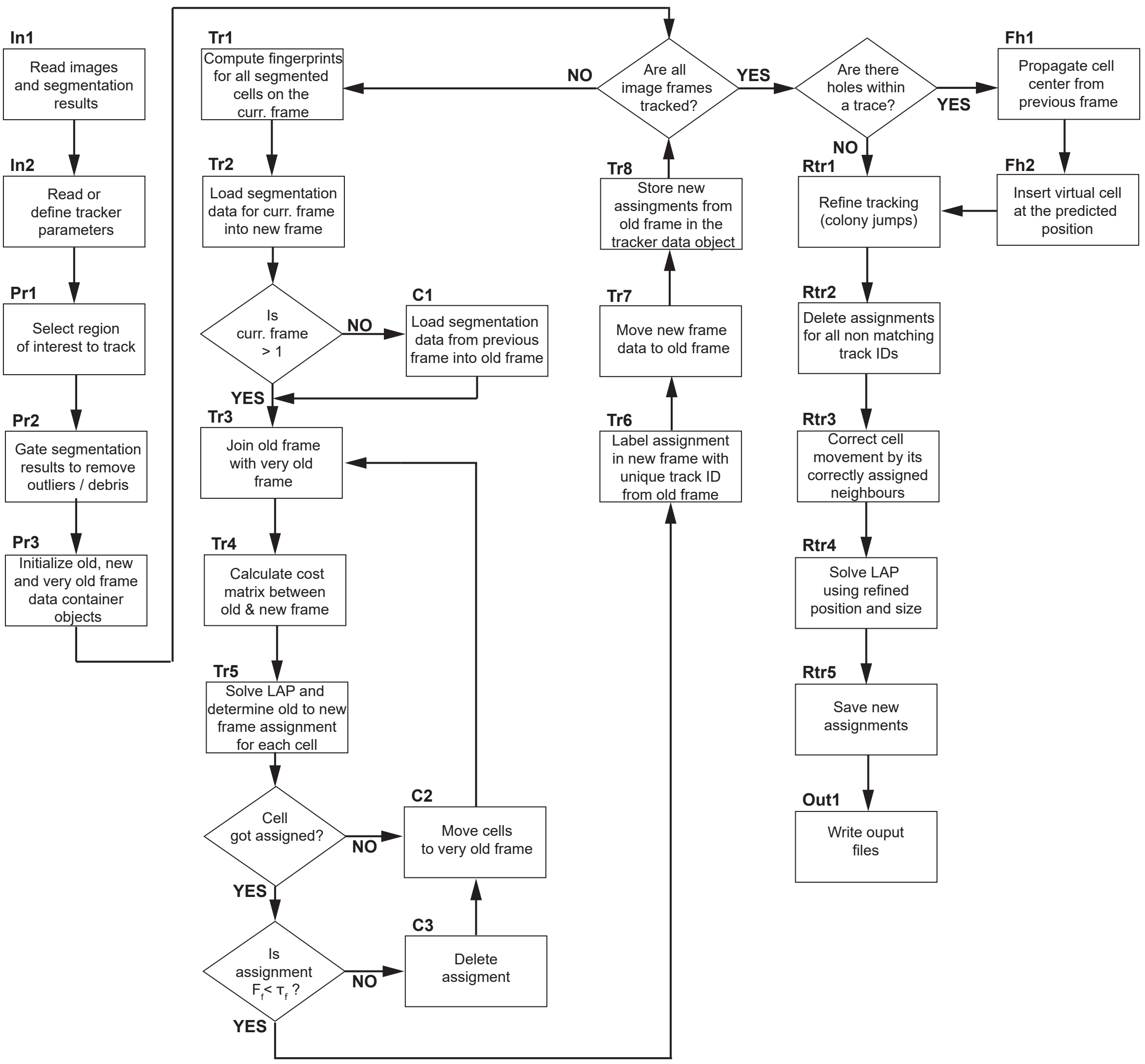


**Figure S8. Detailed flowchart of the tracking core algorithm.** Boxes represent processing tasks, white diamonds decisions, and arrows transitions among them. Labels: In, algorithm inputs; Pr, pre-processing tasks; Tr, core tracking tasks; C, conditions following decisions; Fh, post-processing task of filling trace holes; Rtr, post-processing task of tracking refinement; Out, outputs.

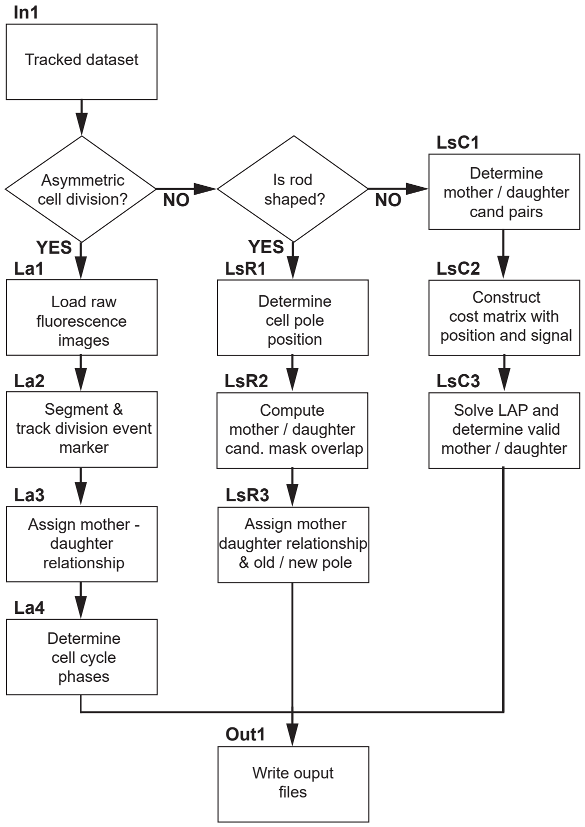


**Figure S9. Flow chart of the lineage reconstruction algorithm**. Lineage reconstruction for asymmetrically dividing cells requires additional division event marker images as input. The mother-daughter assignment is determined based on this marker when it was found repeatably on subsequent frames. For symmetrically dividing cells, the assignment is determined geometrically using the cell poles. Labels: In, algorithm inputs; La, asymmetrical lineage; LsR, symmetrical rod shaped lineage; LsC, symmetrical convex shaped lineage. Out, outputs. For symbols, see **Fig. S7**.

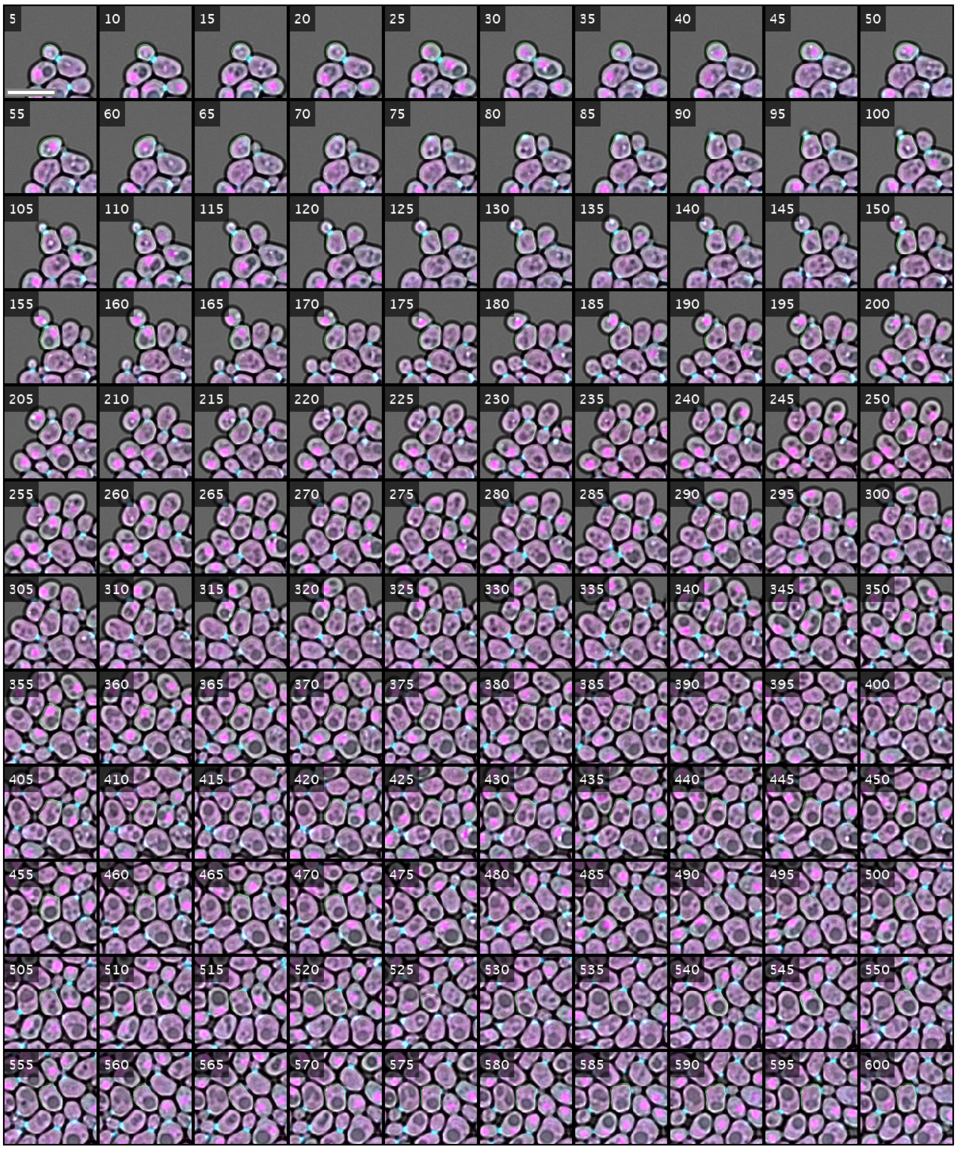


**Figure S10. Cell cycle regulation by Whi5 in *S. cerevisiae* for the selected track in Fig. 6a.** Merge of bright field image channel with the two fluorescent channels depicting Whi5 (magenta) and Myo1 (cyan; see **Methods** for details). Contrast adjusted to brightest pixels for both fluorescent channels. The selected cell from **Fig. 6a** is centered in each tile, with overlay of the outline of its segmentation mask (green). Scale bar = 10 $\mu m$.

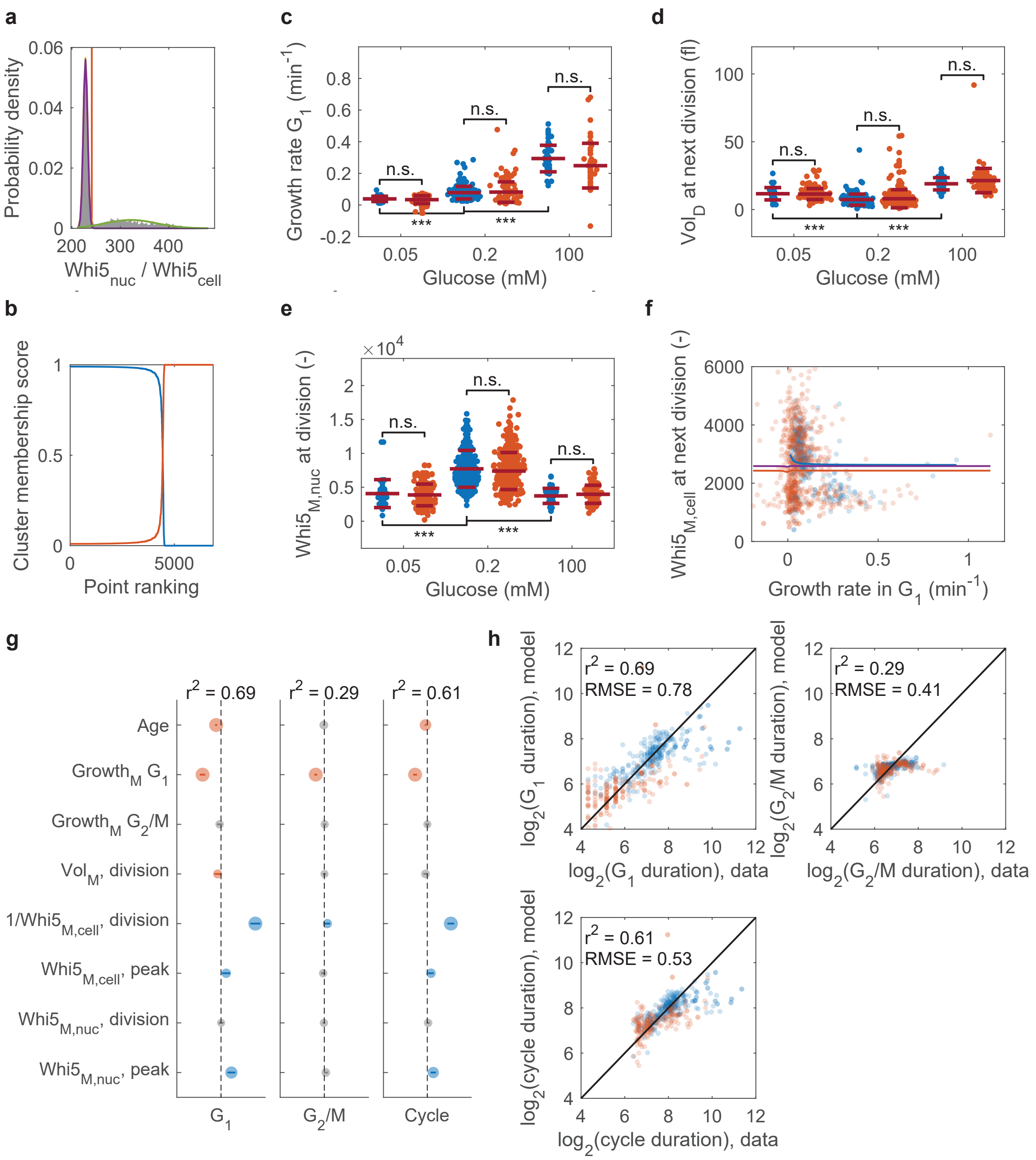


**Figure S11. Cell cycle regulation by Whi5 in *S. cerevisiae*.** **a** Identification of threshold for nuclear concentrations of Whi5 by a two-component Gaussian mixture model (see **Methods** for details). Left: Example of empirical (bars) and fitted (lines) probability densities as well as inferred threshold (red line). Right: Ranking of cluster membership scores based on posterior probability indicates good cluster separation. **b-d** Cell cycle characteristics as a function of cell age and glucose concentration as in **Fig. 6b-e**. **e-g** Correlation plots as in **Fig. 6f-i** with corresponding linear regressions. **h** Volume fraction of daughter cell relative to mother + daughter cell at the time of division. **i** Effects plot for linear models for $G_1$, $G_2$/M, and cell cycle duration (log-scaled response variables; see **Fig. 6j**). **j** Single-cell data vs model predictions for cell cycle duration as in **Fig. 6k,l**.

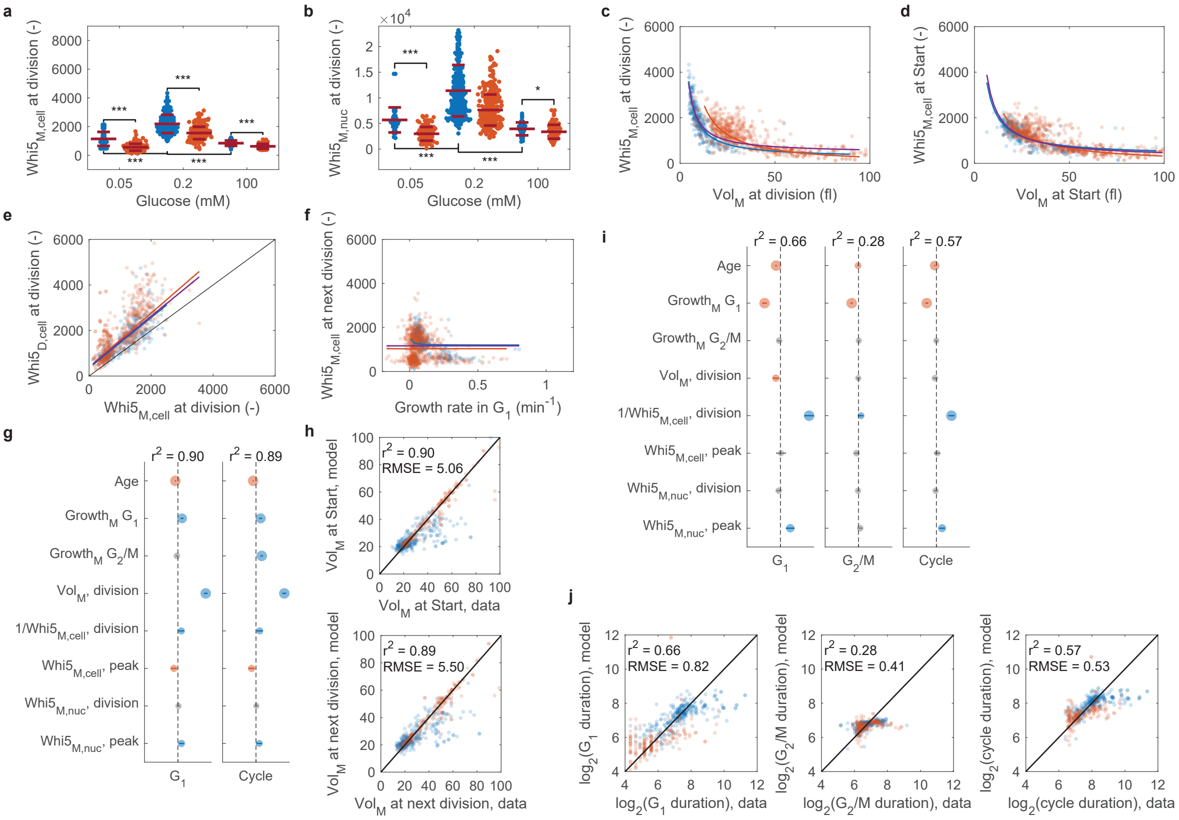


**Figure S12. Whi5 concentration estimation based on cell volume.** Panels correspond to those for concentration estimation based on cell area as follows: **a**: **Fig. 6d**; **b**: **Fig. S11e**; **c**: **Fig. 6f**; **d**: **Fig. 6g**; **e**: **Fig. 6i**; **f**: **Fig. S11f**; **g**: **Fig. 6j**; **h**: **Fig. 6k,l**; **i**: **Fig. S11g**; **j**: **Fig. S11h**.

#### Supplementary Movies

**Figure Movie S1. Lineage results for asymmetric cell division *S. cerevisiae*.** Cells colored by lineage tree of seeding cells from first frame. Details about data set (TTS-SC7) in Table S2

**Figure Movie S2. Lineage results for symmetric cell division (*S. pombe*).** Cells colored by lineage tree of seeding cells from first frame. Details about data set (TTS-SP4) in Table S2

**Figure Movie S3. Lineage results for symmetric cell division (*B. megaterium*).** Cells colored by lineage tree of seeding cells from first frame. Details about data set (TTS-Bmeg) in Table S2

**Figure Movie S4. Lineage results for symmetric cell division HeLa cells.** Subset of TTS-Fluo-N2DL-HeLa. Cells colored by lineage tree of seeding cells from first frame. Details about data set in Table S2

**Figure Movie S5. Lineage results for symmetric cell division HeLa cells.** Full dataset. Cells colored by lineage tree of seeding cells from first frame. Details about data set (TTS-Fluo-N2DL-HeLa) in Table S2

#### Supplementary Tables

**Table S1. Table describing all parameters of *Trac$^X$*.** Parameter; Parameter symbols equal the formulas as described in **Methods**. Parameter names; Parameter names equal the names use din the code of *Trac$^X$*. Unit; SI unit (where applicable). Parameter description; Brief parameter description of what each parameter does.

| Parameter | Parameter name | Unit | Parameter description |
|---|---|---|---|
| $\delta_{max}$ | maxCellCenterDisplacement | px | Max. displacement of segmented objects in pixel allowed in any direction for consecutive image frames (depends on pixel size and magnification). |
| $m_{(s)_{max}}$ | maxExpectedMovement | px | Max expected (linear) movement of one or multiple segmented objects as i.e colony jump in pixel (depends on pixel size and magnification). Default to 2.5· the average cell diameter. Should be adjusted to largest observed movement. |
| $\rho_{\mu}$ | averageCellSizeGrowth | % | Average fractional growth of an identical segmented object between two subsequent image frames (i.e. cell area or nuclear area). |
| $\rho_{dec}$ | maxCellSizeDecrease | % | Max. fractional segmented object size decrease for subsequent image frames. |
| $\rho_{inc}$ | maxCellSizeIncrease | % | Max. fractional segmented object size increase for subsequent image frames. |
| $\theta_{max}$ | maxMajorAxisRotation | deg | Max. tolerated major axis rotation (not needed for round objects). |
| $p$ | individualFunctionPenalty | - | Individual penalty maximum for each cost function (i.e. for position, area, rotation). |
| $\sigma_{max}$ | maxTrackFrameSkipping | frame | Max. number of frames where a segmented object is allowed to be missing from a track. |
| | usedFunctionsForCostMatrix | - | Array specifying which cost functions to consider to build final cost matrix. |
| $\overline{\varnothing(s_{1:n})}$ | meanCellDiameter | px | Mean cell diameter of all segmented objects of all image frames of the whole experiment. |
| | meanCellDiameterScalingFactor | - | Mean cell diameter scaling factor to scale the max allowed linear motion of cells in respect to the mean cell diameter. |
| $f_l$ | fingerprintHalfWindowSideLength | px | Half window side length of a square around the centroid coordinates of a segmented object to be used to for fingerprint computation. |
| $f_r$ | fingerprintResizeFactor | - | Resize factor applied to image matrix before fingerprint calculation. |
| $f_q$ | fingerprintMaxConsideredFrequencies | - | Number of DCT frequencies to include for fingerprint computation. |

*continues on next page*

| | | | |
|---|---|---|---|
| $\tau_f$ | fingerprintDistThreshold | - | Fraction fingerprint threshold $\tau_f$ below which two fingerprints are considered to be identical. |
| $r_{neigh}$ | neighbourhoodSearchRadius | px | Radius around a centroid coordinates of a segmented object used to detect the neighbouring cells. |
| $r_{filt_x}$ | radiusVectorFilterX | px | Radius equals to the first standard deviation of a 2D Gaussian distribution around a cell center in X. The radius is used calculate the neighbourhood weighted cell center displacement in X. |
| $r_{filt_y}$ | radiusVectorFilterY | px | Radius equals to the first standard deviation of a 2D Gaussian distribution around a cell center in Y. The radius is used calculate the neighbourhood weighted cell center displacement in Y. |
| $r_{filt_z}$ | radiusVectorFilterZ | px | Radius equals to the first standard deviation of a 2D Gaussian distribution around a cell center in Z. The radius is used calculate the neighbourhood weighted cell center displacement in Z. |
| | divisionMarkerEdgeSensitivityThresh | - | Edge sensitivity threshold for division marker segmentation. |
| | divisionMarkerConvexAreaUpperThresh | - | Convex area upper threshold used for division marker segmentation selection to exclude segmentation artefacts. |
| | divisionMarkerConvexAreaLowerThresh | - | Convex area lower threshold used for division marker segmentation selection to exclude segmentation artefacts. |
| $\delta_{max}$ | divisionMarkerMaxObjCenterDisplacement | px | Max. displacement of segmented objects in pixel allowed in any direction for consecutive image frames (depends on pixel size and magnification). |
| $m_{(s)_{max}}$ | divisionMarkerMaxExpectedMovement | px | Max expected (linear) movement of one or multiple segmented objects as i.e colony jump in pixel (depends on pixel size and magnification). Default to 2.5$\cdot$ the average cell diameter. Should be adjusted to largest observed movement. |
| $\rho_\mu$ | divisionMarkerAverageObjSizeGrowth | % | Average fractional growth of an identical segmented object between two subsequent image frames (i.e. cell area or nuclear area). |
| $\rho_{dec}$ | divisionMarkerMaxObjSizeDecrease | % | Penalty for maximum fractional segmented object size decrease for subsequent image frames. |
| $\rho_{inc}$ | divisionMarkerMaxObjSizeIncrease | % | Penalty for maximum fractional segmented object size increase for subsequent image frames. |
| $\theta_{max}$ | divisionMarkerMaxMajorAxisRotation | deg | Max. tolerated major axis rotation (not needed for round objects). |
| $p$ | divisionMarkerIndividualFunctionPenalty | - | Individual penalty maximum for each cost function (i.e. for position, area, rotation). |

*continues on next page*

| | | | |
|---|---|---|---|
| $\sigma_{max}$ | divisionMarkerMaxTrackFrameSkipping | Frame | Max. number of frames where a segmented object is allowed to be missing from a track. |
| | divisionMarkerUsedFunctionsForCostMatrix | - | Array specifying which cost functions to consider to build final cost matrix. |
| $\overline{\varnothing(s_{1:n})}$ | divisionMarkerMeanCellDiameter | px | Mean cell diameter of all segmented objects of all image frames of the whole experiment. |
| | divisionMarkerMeanCellDiameterScalingFactor | - | Mean cell diameter scaling factor to scale the max allowed linear motion of cells in respect to the mean cell diameter. |
| $f_l$ | divisionMarkerFingerprintHalfWindowSideLength | px | Half window side length of a square around the centroid coordinates of a segmented object to be used to for fingerprint computation. |
| $f_r$ | divisionMarkerFingerprintResizeFactor | - | Resize factor applied to image matrix before fingerprint calculation. |
| $f_q$ | divisionMarkerFingerprintMaxConsideredFrequencies | - | Number of DCT frequencies to include for fingerprint computation. |
| $\tau_f$ | divisionMarkerFingerprintDistThreshold | - | Fraction fingerprint threshold $\tau_f$ below which two fingerprints are considered to be identical. |
| $r_{neigh}$ | divisionMarkerNeighbourhoodSearchRadius | px | Radius around a centroid coordinates of a segmented object used to detect the neighbouring cells. |
| $r_{filt_x}$ | divisionMarkerRadiusVectorFilterX | px | Radius equals to the first standard deviation of a 2D gaussion distribution around a cell center in X. The radius is used calculate the neighbourhood weighted cell center displacement in X. |
| $r_{filt_y}$ | divisionMarkerRadiusVectorFilterY | px | Radius equals to the first standard deviation of a 2D gaussion distribution around a cell center in Y. The radius is used calculate the neighbourhood weighted cell center displacement in Y. |
| $r_{filt_z}$ | divisionMarkerRadiusVectorFilterZ | px | Radius equals to the first standard deviation of a 2D gaussion distribution around a cell center in Z. The radius is used calculate the neighbourhood weighted cell center displacement in Z. |
| | divisionMarkerLineProfileLength | px | Maximum length of the line profile between two cell centers are re-sampled to. Used to be able to compare the profiles. |
| | divisionMarkerLineProfilePeakLowerBound | px | Lower bound where a peak on line profile between two cell centers is still accepted. Used as threshold. |
| | divisionMarkerLineProfilePeakUpperBound | px | Upper bound where a peak on line profile between two cell centers is still accepted. Used as threshold. |
| | data3D | bool | Is data 2D (0) or 3D (1) |
| | pixelsPerZPlaneInterval | - | The number of pixels that one Z-stack interval is equal to. |
| | active3DFingerprint | bool | Is 3D fingerprint used or closest z stack method |

*continues on next page*

| | | | |
|---|---|---|---|
| $l_{thumb}$ | cellThumbSideLength | px | Thumbnail side length around the center of a segmented object. Used i.e to cut out all cells contained in a single track from all image frames for visualization. |
| $\alpha$ | maskOverlayAlpha | - | Transparency (alpha) level for segmentation mask overlay on raw images for visualization. |
| | debugLevel | - | Parameter defining the debug level. Level 0 shows no output. Level 1 shows general information (info mode). Level 2 shows additional debug info (debug mode). |

**Table S2. Tracker test data sets used in this study.** Abbreviations: BF, brightfield; PhC, phase contrast; FL, fluorescence.

| **Name** | **Species** | **Image modality** | **Image interval (min)** | **Frame count** | **Cell number span** | **Source** |
|---|---|---|---|---|---|---|
| TTS-YIT-TS1 | *S. cerevisiae* | BF | 3 | 60 | 14 - 26 | [2] |
| TTS-YIT-TS2 | *S. cerevisiae* | BF | 3 | 30 | 4 - 6 | [2] |
| TTS-YIT-TS3 | *S. cerevisiae* | BF | 3 | 20 | 101 - 128 | [2] |
| TTS-YIT-TS4 | *S. cerevisiae* | BF | 3 | 20 | 171 - 237 | [2] |
| TTS-YIT-TS5 | *S. cerevisiae* | BF | 3 | 20 | 140 - 173 | [2] |
| TTS-YIT-TS6 | *S. cerevisiae* | BF | 2 | 10 | 36 - 49 | [2] |
| TTS-YIT-TS7 | *S. cerevisiae* | BF | 2 | 10 | 129 - 184 | [2] |
| TTS-YIT-TS8 | *S. cerevisiae* | BF | 3 | 30 | 60 - 88 | [2] |
| TTS-YIT-TS9 | *S. cerevisiae* | PhC | 3 | 30 | 41 - 68 | [2] |
| TTS-YIT-TS10 | *S. cerevisiae* | PhC | 1.5 | 30 | 16 - 16 | [2] |
| TTS-Bmeg | *B. megaterium* | BF | 5.09 | 56 | 8 - 160 | [3] |
| TTS-60mrnaCropped | Bacteria | PhC | ? | 60 | 12 - 28 | [4] |
| TTS-Fluo-N2DH-GOWT1 | mESCs | FL | 5 | 92 | 16 - 17 | [5] |
| TTS-Fluo-N2DL-HeLa | HeLa | FL | 30 | 92 | 4-20/ 42-264 | [5] |
| TTS-SC7 | *S. cerevisiae* | BF | 5 | 115 | 3 - 262 | this study |
| TTS-SC9 | *S. cerevisiae* | BF & FL | 2 | 45 | 553 - 648 | this study |
| TTS-SP4 | *S. pombe* | BF | 2 | 100 | 4 - 111 | [6] |

**Table S3. Primers used in this study.** The primers have been designed to tag the proteins Whi5 and Myo1 according to our Crispr protocol (unpublished; available on request).

| Name | Target | Purpose | Direction | Sequence |
|---|---|---|---|---|
| FRO4253 | Myo1 P5 PacI XFP | Myo1-tagging | Fwd | 5’-AAGCCGTTATGAATCTACCATGATAGACTCGAAAAATATTGATAGTAACAATGCACAGAGTAAAATTTTCAGTGGTGACGGTGCTGGTTTAATTAAC-3’ |
| FRO4254 | Myo1 AscI before term XFP | Myo1-tagging | Rev | 5’-ATAAAGGATATAAAGTCTTCCAAATTTTTAAAAAAAAGTTCGCTTATTTAGAAGTGGCGC-3’ |
| FRO4255 | Myo1 Crispr target seq | Myo1-tagging | Fwd | 5’-GCGCCGGCTGGGCAACACCTTCGGGTGGCGAATGGGACCCGTTATGAATCTACCATGATGTTTTAGAGCTAGAAATAGCAAGTTAAAATAAGGC-3’ |
| FRO4256 | Myo1 Crispr target seq | Myo1-tagging | Rev | 5’-GCCTTATTTTAACTTGCTATTTCTAGCTCTAAAACATCATGGTAGATTCATAACGGGTCCCATTCGCCACCCGAAGGTGTTGCCCAGCCGGCGC-3’ |
| FRO4516 | mKO$\kappa$ end Whi5 term | Whi5-tagging | Rev | 5’-GGGTGCAGCAGCGGGCGCGGCTGCACTAACTCaGAGATTGCGGAGAAAAAACTCGTACTACCACATTAAGAATGAGCAACAGCATCTTCAACTT-3’ |
| FRO4517 | Whi5 Crispr target seq | Whi5-tagging | Fwd | 5’-GCGCCGGCTGGGCAACACCTTCGGGTGGCGAATGGGACCGAGTTTTTTCTCCGCAATCTGTTTTAGAGCTAGAAATAGCAAGTTAAAATAAGGC-3’ |
| FRO4518 | Whi5 Crispr target seq | Whi5-tagging | Rev | 5’-GCCTTATTTTAACTTGCTATTTCTAGCTCTAAAACAGATTGCGGAGAAAAAACTCGGTCCCATTCGCCACCCGAAGGTGTTGCCCAGCCGGCGC-3’ |
| FRO4579 | FRP2008 mKO$\kappa$ | Whi5-tagging | Fwd | 5’-AGCGGGCGCGGCTGCACTAACTCaGAGATTGCGGAGAAAAAACTCGTACTACCACATTAAGAATGAGCAACAGCATCTTCAAC-3’ |

**Table S4. Plasmids used in this study.** Crispr plasmids required for our Crispr protocol (unpublished; available on request).

| Name | Backbone | Description/Insert | Source | Addgene |
|---|---|---|---|---|
| FRP2061 | pCerCC | Crispr plasmid | lab stock | - |
| FRP301 | pFA6a-link-Kan | mKate2(3x) | lab stock | - |
| FRP2008 | pFA6a-link-Kan | mKO$\kappa$ | [6] | #159312 |

**Table S5. Yeast strains used in this study**. All strains have the genetic background FY4. Available on request.

| Name | Description | Genotype | Mating type | Source |
|---|---|---|---|---|
| FRY1398 | BY4700 | MATa & ura3Δ0 | haploid | [7], ATCC® 200866™ |
| FRY1400 | BY4707 | MAT$\alpha$ & met15Δ0 | haploid | [7], ATCC® 200871™ |
| FRY2023 | | FRY1398 & FRY1400 | diploid | lab stock |
| FRY2031 | | FRY2023, MAT$\alpha$ | haploid | lab stock |
| FRY2795 | | FRY2031, Whi5-mKO$\kappa$, Myo1-mKate2(3x) | haploid | this study |
